# Quantifying the RNA cap epitranscriptome reveals novel caps in cellular and viral RNA

**DOI:** 10.1101/683045

**Authors:** Jin Wang, Bing Liang Alvin Chew, Yong Lai, Hongping Dong, Luang Xu, Seetharamsingh Balamkundu, Weiling Maggie Cai, Liang Cui, Chuan Fa Liu, Xin-Yuan Fu, Zhenguo Lin, Pei-Yong Shi, Timothy K. Lu, Dahai Luo, Samie R. Jaffrey, Peter C. Dedon

## Abstract

Chemical modification of transcripts with 5’ caps occurs in all organisms. Here we report a systems-level mass spectrometry-based technique, CapQuant, for quantitative analysis of the cap epitranscriptome in any organism. The method was piloted with 21 canonical caps – m^7^GpppN, m^7^GpppNm, GpppN, GpppNm, and m^2,2,7^GpppG – and 5 “metabolite” caps – NAD, FAD, UDP-Glc, UDP-GlcNAc, and dpCoA. Applying CapQuant to RNA from purified dengue virus, *Escherichia coli*, yeast, mice, and humans, we discovered four new cap structures in humans and mice (FAD, UDP-Glc, UDP-GlcNAc, and m^7^Gpppm^6^A), cell- and tissue-specific variations in cap methylation, and surprisingly high proportions of caps lacking 2’-*O*-methylation, such as m^7^Gpppm^6^A in mammals and m^7^GpppA in dengue virus, and we did not detect cap m^1^A/m^1^Am in humans. CapQuant accurately captured the preference for purine nucleotides at eukaryotic transcription start sites and the correlation between metabolite levels and metabolite caps. The mystery around cap m^1^A/m^1^Am analysis remains unresolved.

## INTRODUCTION

Nearly all forms of RNA are post-transcriptionally modified on the nucleobases or ribose (1), including the 5’-terminal “caps” on messenger (mRNA) and other RNAs (2). The canonical cap on most eukaryotic and viral mRNAs is comprised of *N*^7^-methylguanosine (m^7^G) linked to the first nucleotide of the RNA by a reverse 5’-5’ triphosphate bridge (**Figure 1A**) (2). This m^7^GpppX cap in its various forms (2) is absent in bacterial and archaeal transcripts. In many lower eukaryotes, including yeast, mRNAs contain mainly m^7^GpppN (cap 0), whereas in higher eukaryotes, the 5’ penultimate and antepenultimate nucleotides can be 2’-*O*-methylated to different extents to generate m^7^GpppNm (cap 1) and m^7^GpppNmpNm (cap 2) structures (2). The m^7^GpppX cap has several important biological functions, such as protecting mRNA from degradation by 5’-exoribonucleases, directing pre-mRNA splicing and nuclear mRNA export, facilitating recognition by eukaryotic translation initiation factor 4E, and regulating various aspects of mRNA fate and function, including mRNA stability and mRNA translation (2). In addition, the ribose 2’-*O* methylation (Nm) at the 5’ penultimate nucleotide is thought to be a molecular signature that discriminates self and non-self mRNA, and thus functions in antiviral defense (3).

**Figure 1.**
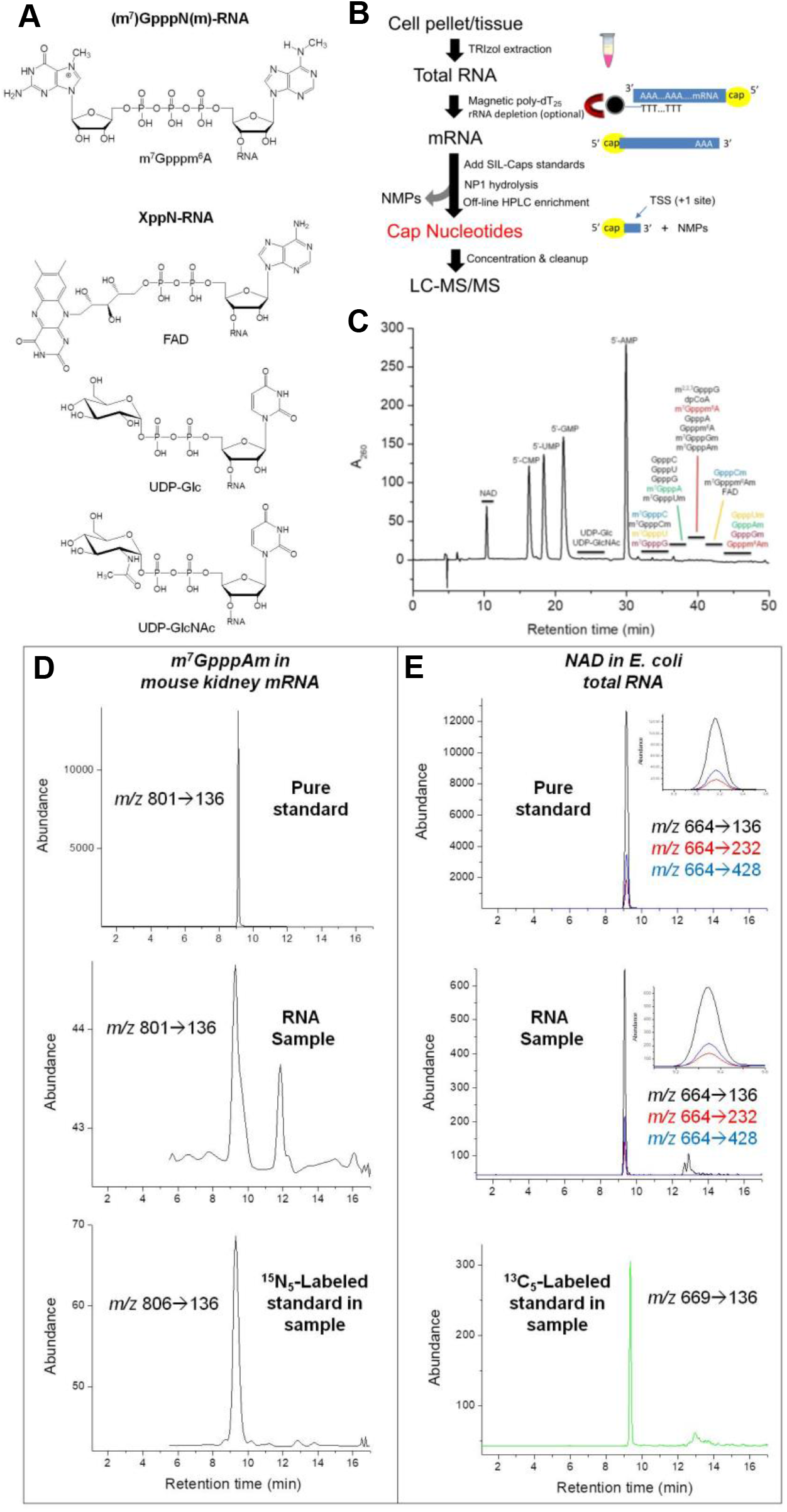
Analysis of 5’ cap structures in RNA by CapQuant. (**A**) Chemical structures of 5’ caps. (**B**) Workflow for CapQuant applied to eukaryotic mRNA. (**C**) A representative HPLC trace for the separation of the enzymatic digestion mixture of RNA. (**D,E**) Illustration of CapQuant for m^7^GpppAm in mRNA from mouse kidney (**D**), and NAD in total RNA from *E. coli* (**E**), showing HPLC elution profiles and MS/MS transitions (*m/z* X→Y) for unlabeled pure standard (top), the RNA sample (middle), and isotope-labeled standard spiked into the RNA sample (bottom). Similar illustrations of CapQuant for all other caps are shown in **Supplementary Figure S8**

The family of eukaryotic RNA caps has recently expanded to include a variety of GpppX variants and non-canonical structures, such as the non-methylated guanosine cap (GpppN) in insect oocyte mRNA (4). Building on the m^7^GpppAm motif, Moss and colleagues showed that up to 30% of caps in animal and viral mRNAs are also methylated at N^6^ of Am (m^6^Am) (5). Multiple methylations also occur on the cap 5’-G, such as di- and tri-methylguanosine caps (e.g., m^2,2,7^GpppN) in viral RNAs (6) and a subset of RNAP II-transcribed cellular RNAs, including small nuclear and nucleolar RNAs, and telomerase RNA (7). Perhaps the simplest methylated cap structure involves γ-phosphate methylation of unprocessed 5’-triphosphate (mPPPN) on small RNAs such as mammalian U6 and 7SK, mouse B2, and plant U3 RNAs (7).

A variety of non-canonical caps involving nucleotide metabolites (**Figure 1A**) have also recently been described (8,9). For example, nicotinamide adenine dinucleotide (NAD) and coenzyme A (CoA) were found as cap-like structures in bacterial small RNAs (10) and the NAD cap was also found in yeast and human mRNA and non-coding RNAs (11). Julius and Yuzenkova expanded the potential repertoire of caps by demonstrating that a variety of nucleotide metabolites could initiate transcription by bacterial RNA polymerase (RNA Pol) *in vitro*, including flavin adenine dinucleotide (FAD), uridine diphosphate glucose (UDP-Glc), and uridine diphosphate *N*-acetylglucosamine (UDP-GlcNAc) (9). They also showed that capping with NAD and UDP analogs by bacterial RNA Pol is promoter-specific and stimulates promoter escape (9), suggesting a role for metabolite caps in regulating gene expression. For example, the NAD cap has been shown to influence RNA stability and turnover, and is a substrate for decapping enzymes (11). However, the lack of sensitive and specific analytical methods has hindered the systematic study of the cap landscape dynamics in cells.

Analysis of RNA cap structures has traditionally relied on radioisotope labeling and enzymatic hydrolysis, followed by thin-layer and other types of chromatography to resolve cap structures (12–14). While sensitive, the radiolabeling approach lacks specificity (12) and has the potential to create cellular toxicity artifacts (15,16). While two-dimensional electrophoresis (14) allows multiple caps analysis, it (i) lacks specificity for identifying intact cap structures, (ii) is limited to NpppN caps, (iii) does not provide absolute quantification, and (iv) is semi-quantitative at best. More recently, methods using high-pressure liquid chromatography (HPLC) with spectroscopic or mass spectrometry-based detection (LC-MS) have been developed (17–22). Though LC-MS provides chemical specificity, existing HPLC and LC-MS methods generally lack sensitivity and are not quantitative.

Here, we report a versatile and sensitive method for transcriptome-wide quantification of RNA caps – CapQuant – that combines off-line HPLC enrichment of cap nucleotides with isotope-dilution, chromatography-coupled triple-quadrupole mass spectrometry (LC-MS/MS) to enable absolute quantification of any type of RNA cap structure with sensitivity (amol-fmol) and chemical specificity. Piloted with 26 different cap structures, this “omic” approach provides important new insights into the landscape of RNA caps in cellular transcriptomes and viruses, and raises questions about current assumptions about cap biology.

## MATERIALS AND METHODS

### Cell and virus culture

CCRF-SB human B lymphoblasts (a gift from Dr. Jianzhu Chen, Singapore-MIT Alliance for Research and Technology) were cultured in RPMI-1640 supplemented with 10% FBS, 50 μg/ml streptomycin and 50 units/ml penicillin at 37 °C and 5% CO_2_. The cells were collected by centrifugation at 350 g for 10 min at 4 °C. *Saccharomyces cerevisiae* strain W1588-4C (a gift from Dr. Graham C. Walker, Massachusetts Institute of Technology) was grown exponentially in YPD medium (1% yeast extract, 2% peptone, 2% glucose) at 30 °C with shaking at 200 rpm. *E. coli* K-12 DH5α cells were grown exponentially in LB broth at 37 °C with shaking (220 rpm) to stationary phase. The cells were collected by centrifugation (4,000 g at 4 °C) and washed once with ice-cold PBS. All cells were stored at −80 °C until total RNA extraction. The preparation and culture of DENV-2 strain TSV01 and isolation of the viral particles were conducted as described previously (23). Briefly, mosquito cells C6/36 were infected with DENV-2 strain TSV01 at an MOI (multiplicity of infection) of 0.1. The infected cells were incubated at 29 °C for 5 days. The virus particles in cell culture supernatant were precipitated by adding 8% PEG8000 (w/v) and incubating the mixture overnight at 4 °C. The precipitated virus particles were then resuspended in NTE buffer (120 mM NaCl, 12 mM Tris-HCl, 1 mM EDTA, pH 8.0) and concentrated by pelleting through a 24% (w/v) sucrose cushion at 75,000 g for 1.5 h at 4 °C. The virus pellet was resuspended into 4% (w/v) potassium tartrate in NTE buffer and centrifuged at 149,000 g for 2 h at 4 °C. The viruses were further purified by ultracentrifugation using a 10–30% (w/v) potassium tartrate gradient. The virus band was collected and concentrated using a 100 kDa centrifugal filter.

### Mouse tissues

Three female C57BL/6 mice were bred in Comparative Medicine, National University of Singapore (NUS), following the polices and guidelines of the NUS Institutional Animal Care and Use Committee. The mice were sacrificed at 4-6 months of age for collection of tissues, which were snap-frozen in liquid nitrogen and stored at −80 °C.

### Cap nucleotide standards

GpppA, GpppG, m^7^GpppA and m^7^GpppG were purchased from New England Biolabs (NEB; Ipswich, MA USA). NAD, FAD, UDP-Glc, UDP-GlcNAc and dpCoA were purchased from Sigma Chemical Co. (St. Louis, MO USA). m^2,2,7^GpppG was purchased from Jena Bioscience (Jena, Thuringia, Germany). [^13^C_5_]-β-Nicotinamide adenine dinucleotide ammonium salt (^13^C_5_-NAD) and [^13^C_5_]-flavin adenine dinucleotide ammonium salt hydrate (^13^C_5_-FAD) were purchased from Medical Isotopes (Pelham, NH USA). [^13^C_6_]-Uridine diphosphate glucose (^13^C_6_-UDP-Glc) disodium salt and uridine diphosphate *N*-acetylglucosamine-^13^C_6_ (^13^C_6_-UDP-GlcNAc) disodium salt were from Omicron Biochemicals (South bend, IN USA). GpppAm- and m^7^GpppAm-capped RNA oligos were synthesized by *in vitro* 2’-*O*-methylation of the penultimate adenosine residue of G-capped and m^7^G-capped dengue RNA representing the first 211 nucleotides of DENV-4 genome (strain MY-22713), respectively, by ScriptCap 2’-*O*-Methyltransfease. The dengue RNA was *in vitro* transcribed from PCR products amplified using an infectious cDNA clone as a template and the pairs of primer as below. Forward primer: 5’-CAGTAATACGACTCACTATTAGTTGTTAGTCTGTGTGGAC-3’, reverse primer: 5’-TAGCACCATCCGTAAGGGTC-3’. G-capped and m^7^G-capped RNA were generated using MEGAshortscript T7 Transcription Kit (Invitrogen) according to the manufacturer’s instructions. Briefly, NTPs (ATP =6 mM, GTP = 7.5 mM, CTP = 7.5 mM, UTP = 7.5 mM) and GpppA (1.5 mM) or m^7^GpppA (1.5 mM) were added into the reaction. Capped RNA was purified by passing through two G-25 size columns (GE Healthcare), extracted with phenol–chloroform, and precipitated with ethanol. The purified capped RNA was subjected to 2’-*O* methylation using ScriptCapTM 2’-*O*-Methyltransferase (Epicentre) in the presence of cold S-adenosyl methionine (SAM) following the Instruction Manual. The methylated RNA oligos were purified in the same fashion as the capped RNA. RNA oligos (22 nt) with the following caps were synthesized by *in vitro* reaction of pppXGGCUCGAACUUAAUGAUGACG (Bio-Synthesis Inc., X = C, U, G, A, m^6^A, Cm, Um or Gm) with the Vaccinia Capping System (VCS) in the presence or absence of SAM, according to manufacturer directions: GpppC, GpppU, Gpppm^6^A, m^7^GpppC, m^7^GpppU, m^7^Gpppm^6^A, GpppCm, GpppUm, GpppGm, m^7^GpppCm, m^7^GpppUm and m^7^GpppGm. 500-1000 pmol of each pppXGGCUCGAACUUAAUGAUGACG RNA oligo was heated at 65 °C for 5 min and then chilled on ice for 5 min. To the RNA was then added 10 μl of 10× Capping Buffer (NEB), 5 μl of 10 mM GTP, VCS (NEB, 50 U every two hours) and water, making a final volume of ~100 μl. For the synthesis of m^7^GpppN and m^7^GpppNm, 20 mM of cold SAM (2 μl per hour) was also added. The mixture was briefly mixed by vortexing and then incubated at 37 °C for 4 h, with the enzyme subsequently removed by extraction with chloroform:isoamyl alcohol 24:1 (Sevag, Fluka). The RNA in the aqueous layer was then purified by passing through a 3000 Da spin filter, followed by washing three times with water. Gpppm^6^Am- and m^7^Gpppm^6^Am-capped RNA oligos were synthesized and purified as described previously (24). The synthesis and purification of RNA oligos with [^15^N_5_]-labeled G or m^7^G in the cap (GpppN, N = C, U, G, A or m^6^A; m^7^GpppN, N = C, U, G, A or m^6^A; GpppNm, Nm = Cm, Um, Gm or Am; and m^7^GpppNm, Nm = Cm, Um, Gm or Am) were conducted with 200-500 pmol of each pppXGGCUCGAACUUAAUGAUGACG oligo as RNA substrate in the same fashion except that [^15^N_5_]-GTP (Sigma Chemical Co.) was used instead of GTP in the VCS reaction step. RNA oligo carrying a [^15^N_5_]-m^7^Gpppm^6^Am cap was synthesized as follows. Briefly, 500 pmol of RNA oligo pppm^6^AGGCUCGAACUUAAUGAUGACG (Bio-Synthesis Inc.; Lewisville, TX USA) was heated at 65 °C for 5 min and then chilled on ice for 5 min. To the RNA was then added 5 μl of 10× Capping Buffer, 5 μl of 10 mM GTP, 20 mM of cold SAM (2 μl per hour), VCS (20 U every 2 h), vaccinia mRNA 2’-*O*-methyltransferase (NEB, 250 U every 2 h) and water, making a final volume of ~50 μl. The mixture was briefly mixed by vortexing and then incubated at 37 °C for 4 h, with the enzymes subsequently removed by extraction with Sevag. The RNA in the aqueous layer was then purified in the same way as described above. RNA oligo carrying a [^15^N_5_]-Gpppm^6^Am cap was synthesized as follows. Briefly, 250 pmol of oligo pppm^6^AGGCUCGAACUUAAUGAUGACG (Bio-Synthesis Inc.; Lewisville, TX USA) was heated at 65 °C for 5 min and then chilled on ice for 5 min. To the RNA was then added 5 μl of 10× Capping Buffer, 2.5 μl of 10 mM [^15^N_5_]-GTP, 20 mM of cold SAM (1 μl per hour), VCS (10 U every two hours) and water, making a final volume of ~50 μl. The mixture was briefly mixed by vortexing and then incubated at 37 °C for 4 h, with the enzyme subsequently removed by extraction with Sevag. The RNA in the aqueous layer was purified in the same way as described above. The purified RNA was heated at 65 °C for 5 min and then chilled on ice for 5 min. To the RNA was then added 10 μl of 400 mM Tris-HCl pH 7.5, 10 μl of 50 mM DTT, 20 mM of cold SAM (2 μl per hour), DENV NS5 methyltransferase (200 pmol every two hours) and water, making a final volume of ~100 μl. The mixture was briefly mixed by vortexing and then incubated at 37 °C for 4 h, with the enzyme subsequently removed by extraction with Sevag. The RNA in the aqueous layer was purified in the same way as described above. All synthetic capped oligos were digested with NP1 (30 mM sodium acetate pH 5.5 and 1 mM ZnCl_2_, 37 °C) and the caps purified by ion-pairing HPLC, with cap fractions concentrated and cleaned up by Speed-vac, as described in the HPLC section below. All purified synthetic cap dinucleotides were >99% or >98% pure based on HPLC and were characterized by high-resolution mass spectrometry (HRMS) (**Supplementary Table S2**) and MS/MS analyses (**Supplementary Figure S1**). The synthesis of RNA oligo containing a mixture of m^7^Gpppm^1^A and m^7^Gpppm^1^Am in the 5’ cap and the release and purification of m^7^Gpppm^1^A and m^7^Gpppm^1^Am were conducted in the same fashion. The purified m^7^Gpppm^1^A and m^7^Gpppm^1^Am were >98% and >99% pure respectively based on HPLC, with their identity confirmed by MS/MS analysis (**Supplementary Figure S1**) and successful detection of m^1^A and m^1^Am, but not m^6^A and m^6^Am, respectively by LC-MS/MS (**Supplementary Figure S2**) using the same method as the LC-MS/MS method described below for Dimroth rearrangement analysis following hydrolysis into nucleosides by RNA 5’ pyrophosphohydrolase (RppH, NEB) and shrimp alkaline phosphatase (SAP, NEB). The concentrations of the caps, m^7^Gpppm^1^A and m^7^Gpppm^1^Am were measured by their UV absorbance at 260 nm. The isotopic purity of the caps was found to be better than 99.6% (data not shown) based on LC-MS/MS analyses.

### m^1^A, m^6^A, m^1^Am and m^6^Am nucleoside standards

m^1^A, m^6^A and m^6^Am were purchased from Berry and Associates (Dexter, MI USA). m^1^Am was synthesized by reaction of methyl iodide (0.3 mL) with 2’-*O*-methyladenosine (100 mg) in anhydrous DMF (2.0 mL) in a closed flask with stirring at ambient temperature for 18 h. The reaction mixture was evaporated under vacuum and triturated with diethyl ether to afford a white solid (120 mg). A portion of this crude solid (40 mg) was dissolved in 3.0 mL of methanol and treated with aqueous ammonia (3.0 mL) by stirring at ambient temperature for 10 min. Following evaporation of solvent under vacuum, the mixture was resolved by chromatography on 200-400 mesh silica gel eluted with 15-20% methanol in dichloromethane with 1% aqueous ammonia to afford m^1^Am (25 mg, 59%) as a white solid. The product was characterized by ^1^H and ^13^C NMR (**Supplementary Figure S3**) and HRMS: ^1^H NMR (DMSO-D_6_, 400 MHz) δ 8.18 (s, 1H), 8.09 (s, 1H), 7.03 (bs, H,), 5.87 (d, *J* = 6.00 Hz, 1H), 5.25 (d, *J* = 5.24 Hz, 1H), 5.14 (t, *J* = 5.54 Hz, 1H), 4.29 (m, 1H), 4.25 (m, 1H), 3.95 (q, *J* = 10.68 Hz, 1H), 3.65 (m, 1H), 3.56 (m, 1H), 3.43 (s, 3H), 3.31 (s, 3H); ^13^C (DMSO-D_6_, 100 MHz) δ 154.8, 149.1, 141.9, 138.1, 123.1, 86.7, 85.9, 83.4, 69.1, 61.8, 58.0, 35.1; HRMS (ESI, *m/z*) calculated for C_12_H_18_N_5_O_4_ [M + H]^+^: 296.1359, found: 296.1370, mass error < 5 ppm.

### H_2_O_2_ and MMS treatment

Treatment of *S. cerevisiae* W1588-4C cells with 6 mM of MMS or 2 mM of H_2_O_2_ was started when the O.D. reached ~0.5. After 1 h treatment, the cells were collected by centrifugation (4,500 g at 4 °C) and washed twice with ice-cold PBS.

### RNA extraction

The total RNA from CCRF-SB pellets was directly extracted with TRIzol reagent (Life Technologies), according to the manufacturer’s protocol. For mice, the liver and kidney tissues were ground under liquid nitrogen into fine powders in a mortar, the total RNA of which were then extracted with TRIzol reagent as described earlier. For yeast, total RNA was extracted with a MasterPure Yeast RNA Purification kit (Epicentre) following the manufacturer’s protocol. For *E. coli*, lysis was performed with lysozyme, before total RNA was extracted with TRIzol reagent as described earlier. Briefly, 0.8 ml of TE buffer (pH 8.0) containing 80 mg lysozyme (Fluka) was added to approximately 3.7 × 10^10^ *E. coli* DH5α cells and the mixture was incubated for 2 h at room temperature. To the mixture was then added 0.6 ml of TE buffer (pH 8.0) containing 60 mg lysozyme, followed by incubation for another 2 h at room temperature. Total RNA was subsequently extracted with TRIzol following the manufacturer’s instructions. The genomic RNA from purified dengue virions was extracted with TRIzol and purified by size-exclusion chromatography as described previously (23,25). The poly(A)-tailed RNA in human CCRF-SB cells was isolated from the total RNA using a Fasttrack MAG Maxi mRNA isolation kit (Life Technologies), whereas the poly(A)-tailed RNA in yeast cells and mouse tissues was isolated from the total RNA using a Dynabeads mRNA Purification kit (Life Technologies) following the manufacturer’s protocols. rRNA depletion of the poly(A)-tailed RNA isolated from yeast cells and mouse tissues was subsequently performed using a GeneRead rRNA Depletion kit (Qiagen), according to the manufacturer’s protocol. The rRNA-depleted RNA was then cleaned up using a RNeasy MinElute Cleanup kit (Qiagen), following the manufacturer’s protocol. No rRNA depletion and subsequent clean-up was performed for the poly(A)-tailed RNA isolated from human CCRF-SB cells because there was no sign of significant rRNA contamination (**Supplementary Figure S4**). All RNA samples were stored at −80 °C before use. The quality of the total RNA (**Supplementary Figure S5**), poly(A)-tailed RNA (**Supplementary Figure S4**), and purified DENV-2 RNA genome (**Supplementary Figure S4**) was assessed using an Agilent Bioanalyzer (Agilent Technologies) with RNA 6000 Nano or Pico chips.

### RNA hydrolysis

Isolated RNA (200 μg for total RNA and 0.6-7.8 μg for mRNA and RNA genome) was incubated with NP1 (1 unit/μg RNA, Sigma) in a solution containing 30 mM sodium acetate pH 5.5, 1 mM ZnCl_2_ and 24 SIL-CNs at 37 °C for 1 h. These SIL-CNs included 200 fmol of NAD, 200 fmol of FAD, 500 fmol of UDP-Glc, 500 fmol of UDP-GlcNAc, 500 fmol of GpppC, 200 fmol of GpppU, 400 fmol of GpppG, 500 fmol of GpppA, 500 fmol of Gpppm^6^A, 500 fmol of m^7^GpppC, 200 fmol of m^7^GpppU, 1000 fmol of m^7^GpppG, 500 fmol of m^7^GpppA, 100 fmol of m^7^Gpppm^6^A, 1000 fmol of GpppCm, 200 fmol of GpppUm, 1000 fmol of GpppGm, 500 fmol of GpppAm, 100 fmol of Gpppm^6^Am, 500 fmol of m^7^GpppCm, 200 fmol of m^7^GpppUm, 500 fmol of m^7^GpppGm, 500 fmol of m^7^GpppAm, and 200 fmol of m^7^Gpppm^6^Am. The enzyme was subsequently removed by extraction with Sevag. The resulting aqueous layer was subjected to off-line HPLC separation for the enrichment of the CNs and their analogs (m^7^Gpppm^1^A and m^7^Gpppm^1^Am) under study.

### HPLC

A 4.6 mm×250 mm Alltima HP C18 column (5 μm in particle size, Hichrom) was used for the enrichment of CNs and their analogs from the enzymatic digestion products of RNA. A solution of 10 mM dibutylammonium acetate (DBAA) in 5% ACN-95% H_2_O (solution A) and 10 mM DBAA in 84% ACN-16% H_2_O (solution B) were used as mobile phases, and the flow rate was 0.8 mL/min. A gradient of 20 min 0% B and 40 min 0-40% B was employed. A typical HPLC trace is depicted in **Figure 1C**. The HPLC fractions eluting approximately at 10.0-12.0,13.5-15.9, 19.0-20.6, 23.0-28.0, 32.0-36.0, 36.0-37.5, 37.5-39.0, 39.0-41.5, 41.5-43.0, and 43.0-46.5 min were pooled for NAD, m^7^Gpppm^1^A, m^7^Gpppm^1^Am, (UDP-Glc and UDP-GlcNAc), (m^7^GpppC, m^7^GpppU, m^7^GpppG and m^7^GpppCm), (GpppC, GpppU, GpppG, m^7^GpppA and m^7^GpppUm), (GpppA, Gpppm^6^A, m^7^Gpppm^6^A, m^7^GpppGm, m^7^GpppAm, m^2,2,7^GpppG and dpCoA), (FAD, GpppCm and m^7^Gpppm^6^Am) and (GpppUm, GpppAm, GpppGm and Gpppm^6^Am), respectively. The collected fractions were dried in the Speed-vac, reconstituted in acetonitrile:water 3:7 (v/v) and dried for three cycles to remove the ion-paring reagent present in the fractions, reconstituted in 8 mM ammonium bicarbonate pH 7.0 (solution C), and injected for LC-MS/MS analysis.

### LC-MS/MS analysis of cap nucleotides

Using purchased and synthetic standards, we defined the HPLC retention times for the 26 CNs and two analogs of them (m^7^Gpppm^1^A and m^7^Gpppm^1^Am) on a Luna Omega PS C18 column (100 × 2.1 mm, 1.6 μm) coupled to an Agilent 1290 HPLC system and an Agilent 6460 triple quad mass spectrometer. The elution was conducted at 15 °C and a flow rate of 200 μL/min, with a gradient of 100% solution C and 0% solution D (methanol) for 5 min, followed by 0% to 48% solution D over a period of 12 min. The HPLC column was coupled to an Agilent 6460 Triple Quad mass spectrometer with an electrospray ionization source in positive or negative mode with the following parameters: gas temperature, 350 °C; gas flow, 11 L/min; nebulizer, 20 psi; sheath gas temperature, 300 °C; sheath gas flow, 12 L/min; capillary voltage, 1,800 V; nozzle voltage, 2,000 V; fragmentor voltage, 135 V; ΔEMV, 400 V. MRM mode was used for detection of product ions derived from the precursor ions for all the 26 unlabeled CNs and 24 SIL-CNs with instrument parameters which mainly included the collision energy (CE) optimized for maximal sensitivity for the CNs (mode, retention time in min, precursor ion of unlabeled CN *m/z*, product ion(s) of unlabeled CN *m/z* (CE), precursor ion of labeled CN *m/z*, product ion of labeled CN *m/z* (CE)): NAD, positive, 9.3, 664, 136 (39 V), 232 (24 V), 428 (30 V), 669, 136 (39 V); FAD, positive, 14.0, 787, 348 (20 V), 136 (44 V), 439 (28 V), 782, 353 (20 V); UDP-Glc, negative, 1.3, 565, 323 (24 V), 79 (76 V), 211 (32 V), 570, 323 (24 V); UDP-GlcNAc, negative, 1.4, 606, 385 (28 V), 273 (36 V), 282 (36 V), 612, 385 (28 V); GpppC, positive, 1.7, 749, 152 (60 V), 754, 157 (60 V); GpppU, positive, 2.0, 750, 152 (28 V), 755, 157 (28 V); GpppG, positive, 2.2, 789, 152 (60 V), 794, 157 (60 V); GpppA, positive, 3.9, 773, 136 (56 V), 778, 136 (56 V); Gpppm^6^A, positive, 8.8, 787, 150 (80 V), 792, 150 (80 V); m^7^GpppC, positive, 1.8, 763, 166 (56 V), 768, 171 (56 V); m^7^GpppU, positive, 1.8, 764, 166 (36 V), 769, 171 (36 V); m^7^GpppG, positive, 5.4, 803, 248 (32 V), 808, 248 (32 V); m^7^GpppA, positive, 10.8, 787, 136 (68 V), 792, 136 (68 V); m^7^Gpppm^6^A, positive, 9.3, 801, 150 (80 V), 806, 150 (80 V); GpppCm, positive, 2.3, 763, 111 (52 V), 768, 111 (52 V); GpppUm, positive, 3.7, 764, 152 (40 V), 769, 157 (40 V); GpppGm, positive, 8.2, 803, 111 (56 V), 808, 111 (56 V); GpppAm, positive, 8.8, 787, 136 (60 V), 792, 136 (60 V); Gpppm^6^Am, positive, 10.2, 801, 150 (72 V), 806, 150 (72 V); m^7^GpppCm, positive, 3.4, 777, 166 (52 V), 782, 171 (52 V); m^7^GpppUm, positive, 6.2, 778, 166 (32 V), 783, 171 (32 V); m^7^GpppGm, positive, 8.5, 817, 166 (68 V), 822, 171 (68 V); m^7^GpppAm, positive, 9.7, 801, 136 (68 V), 806, 136 (68 V); m^7^Gpppm^6^Am, positive, 10.8, 815, 150 (76 V), 820, 150 (76 V); dpCoA, positive, 11.7, 689, 261 (24 V), 348 (20 V), 136 (40 V); m^2,2,7^GpppG, positive, 8.5, 831, 194 (64 V), 248 (28 V), 566 (32 V); m^7^Gpppm^1^A, positive, 4.2, 401, 166 (16 V), 150 (36 V); m^7^Gpppm^1^Am, positive, 9.3, 408, 166 (16 V), 150 (32 V), 111 (36 V).

### Genome-wide nucleotide distribution of TSS

To cross-validate the CapQuant results obtained in this study, transcriptional start site (TSS) nucleotide identities were mined from the 5’ terminal positions of capped transcripts mapped using cap-analysis gene expression (CAGE) approach (26,27). CAGE datasets were chosen over others, such as serial analysis of gene expression, as the CAGE method captures mRNA transcripts at the 7-methylguanosine cap to pulldown the 5’-cDNAs reversely transcribed from them (28) for subsequent tagging and high-throughput sequencing. It achieves genome-wide 1bp-resolution map of TSSs and expression levels. Mapped TSS reads are represented as units of peaks due to varying spread of positions which have first base signals within a promoter, and a reading of greater than 10 read counts and 1 tag per million (TPM) signifies a robust TSS signal. The TSS analysis workflow herein is outlined in **Supplementary Figure S6a**. CAGE data in .bedgraph format for *Sacharromyces cerevisiase BY4741* was obtained from the YeasTSS Atlas (Yeast Transcription Start Site Atlas) (29). While CAGE data for human and mouse was obtained from the FANTOM5 project (Functional ANnoTation Of Mammalian genomes) via http://fantom.gsc.riken.jp/5/datafiles/reprocessed/ (30,31). These datasets were uploaded into the main public Galaxy server (32) into separate history list with the referent genome set to the latest assembly for further processing. First, non-robust TSS signals were removed in yeast data (c4 of .bedgraph file), a score of >1 and <-1 was Filtered for the positive and negative strand respectively. Second, GetFastaBed under BedTools (33) was used to extract the respective TSS nucleotide information in tab-delimited format and force strandedness applied to reverse complement negative sense strand. GetFastaBed for human and mouse data were obtained from thickStart and thickEnd (c7 and c8) positions, Trimmed up to position 1 to obtain the 5’ terminal nucleotide only, Change Case to upper case. Third, Count under Statistics to obtain the TSS nucleotide distribution histograms for human (**Supplementary Figure S6b**), mouse (**Supplementary Figure S6c**). and yeast data (**Supplementary Figure S6d**). As the number of transcripts generated from different TSSs can be very different, the weighted and unweighted nucleotide frequency of TSS could affect correlation accuracy. To account for the weight of TSS usage frequency according to transcript abundance, Datamash was performed by grouping the nucleotides together and summing the CTSS read counts (c5 of.bed file) to obtain the weighted values for human (**Supplementary Figure S6b**), mouse (**Supplementary Figure S6c**) and yeast (**Supplementary Figure S6d**). The work histories can be accessed via https://usegalaxy.org/histories/list_published?f-username=alvin_chew

### Dimroth rearrangement

Due to the limited quantities of m^7^Gpppm^1^A and m^7^Gpppm^1^Am we obtained, we performed the testing of the Dimroth rearrangement with purchased m^1^A and synthetic m^1^Am nucleoside standards (**Supplementary Figure S7**). Because the CCRF-SB mRNA samples were the most abundant mammalian mRNA samples we had and they were the only mRNA samples for which no further purification by rRNA depletion was performed, we chose to use the CCRF-SB mRNA samples for the analysis. We treated a mixture of m^1^A and m^1^Am in the same fashion as CCRF-SB cells or the isolated RNA as we went through the RNA extraction, purification, cleanup and enzymatic digestion steps (**Supplementary Figure S7a**) as described above. The m^1^A, m^6^A, m^1^Am and m^6^Am in the samples were separated on a Hypersil GOLD aQ C18 column (100 × 1 mm, 1.9 μm) coupled to an Agilent 1290 HPLC system and an Agilent 6460 triple quad mass spectrometer. The elution was conducted at 24 °C and a flow rate of 100 μL/min, with a gradient of 100% solution E (0.1% formic acid in water) to 89% solution E-11% solution F (0.1% formic acid in acetonitrile) over a period of 11 min, followed by a gradient of 11% to 80% solution F over a period of 3 min. The HPLC column was coupled to an Agilent 6460 Triple Quad mass spectrometer with an electrospray ionization source in positive mode with the following parameters: gas temperature, 300 °C; gas flow, 5 L/min; nebulizer, 45 psi; sheath gas temperature, 200 °C; sheath gas flow, 5 L/min; capillary voltage, 3,500 V; nozzle voltage, 500 V; fragmentor voltage, 110 V; ΔEMV, 800 V. MRM mode was used for detection of product ions derived from the precursor ions for m^1^A, m^6^A, m^1^Am and m^6^Am with the following instrument parameters (retention time in min, precursor ion *m/z*, product ion *m/z*, CE): m^1^A, 2.4, 282, 150, 15 V; m^6^A, 6.1, 282, 150, 15 V; m^1^Am, 4.5, 296, 150, 15 V; m^6^Am, 7.8, 296, 150, 15 V.

#### RT-qPCR

Quantitative real-time PCR (RT-qPCR) was performed to assess the relative mRNA abundance of a selection of RNA cap modification enzymes including PCIF1 (the enzyme responsible for the synthesis of m^6^Am in mRNA caps), FTO (an RNA *N*^6^-methyladenine demethylase that can act on cap m^6^A/m^6^Am in mammals), DCP2 (a major RNA decapping enzyme in mammals) and CMTR1 (cap 1 2’-*O*-ribose methyltransferase), as well as ALKBH5 (another RNA *N*^6^-methyladenine demethylase) in the total RNA from CCRF-SB cells and mouse liver and kidney tissues. 1 μg of total RNA was reverse transcribed using iScript™ cDNA Synthesis Kit (Bio-Rad) according to the manufacturer’s instructions. The cDNA was subjected to qPCR analysis using BlitzAmp qPCR Master Mix (MiRXES Pte Ltd) according to the manufacturer’s fast thermal cycling instructions on a CFX96 Realtime-PCR System (Bio-Rad). Experiments were performed with three biological and two technical replicates in hard-shell thin wall PCR plates (#HSP9601; Bio-Rad). No template and no reverse transcriptase controls were used to assess primer dimerization and genomic DNA contamination, respectively. Relative gene expression was calculated using a modified comparative method for geometric averaging of two reference genes, Gapdh and Polr2a, for more reliable normalization (34). Data visualization and Student’s t-test statistical analysis was performed using Graphpad Prism software (version 8.0). Error bars represent mean ± s.d., and n.s. means not significant.

## RESULTS

### Development of CapQuant

The workflow for CapQuant (**Figure 1B**) uses nuclease P1 (NP1) to hydrolyze RNA to nucleoside monophosphates (NMPs) while sparing di- and tri-phosphate linkages that characterize the NpppN and NppN caps (24,35). Following removal of NP1, cap structures and 5’-NMPs in the limit digest are resolved by reversed-phase ion-paring HPLC (**Figure 1C**) and cap-containing fractions isolated for subsequent LC-MS/MS quantification. Here we targeted 26 caps that embraced a variety of known and possible structures: m^7^GpppN, m^7^GpppNm, GpppN, GpppNm (N = C, U, G, A or m^6^A), and NAD, FAD, UDP-Glc, UDP-GlcNAc, m^2,2,7^GpppG and dpCoA. The 26 caps were well resolved from 5’-NMPs (**Figure 1C**), separating each member of four isobaric pairs using mobile phases containing the volatile ion-pairing agent dibutylammonium acetate (DBAA). Cap-containing fractions were collected and the volatile ion-pairing agent completely removed by three cycles of drying and reconstitution in acetonitrile:water 3:7 (v/v). Samples were finally reconstituted in ammonium bicarbonate buffer (pH 7.0) for subsequent analysis.

Individual caps were next quantified by isotope-dilution LC-MS/MS, the most rigorous approach for sensitivity, specificity, and quantitative accuracy. HPLC conditions for the LC-MS/MS analysis were systematically optimized using standards for the 26 caps, with assessment of different solid phases (C18/NH_2_ reversed-phase, HILIC, porous graphite), pH values (2.7-9.0), and column temperatures (10-45 °C). The best overall resolution and sensitivity were obtained with a positive-surface C18 column at 15 °C with volatile ammonium bicarbonate (pH 7.0) as a mobile phase. Isotope-labeled standards for 24 of the 26 caps were spiked into RNA samples prior to NP1 hydrolysis and each cap was identified by HPLC retention time and collision-induced dissociation (CID) patterns, using MS parameters optimized for each cap (**Figure 1D and E; Supplementary Figure S8**). Quantification was achieved using a calibration curve for each cap (**Supplementary Figure S9**) generated by multiple-reaction monitoring (MRM), with one MRM transition for m^7^GpppN, m^7^GpppNm, GpppN and GpppNm caps and three MRM transitions for the other 6 caps (**Figure 1D and E, Supplementary Figure S8**). This resulted in limits of detection (LODs) ranging from 19 amol to 13 fmol for 23 caps, and up to 160 fmol for 3 caps (GpppC, GpppCm and GpppGm; **Supplementary Table S1**). As shown in **Figure 1D and E**, which depicts applications of the method to mouse (C57BL/6) kidney mRNA and *Escherichia coli* DH5α total RNA, CapQuant proved to be sensitive, precise, and accurate.

Using this new method, control experiments were performed to ensure complete cap release and stability during sample processing. To confirm that all detected caps were indeed covalently linked to mRNA prior to NP1 digestion and not present as contaminants, we used the method to analyze *S. cerevisiae* mRNA and *E. coli* total RNA except that NP1 was removed from its stock solution with a 3000 Da filter and the filtrate used in the RNA digestion reaction. None of the cap analytes were detectable in subsequent LC-MS/MS analyses, from which we conclude that CapQuant analytes are truly RNA caps. To validate complete and unbiased release of all m^7^G caps from RNA, we quantified release of m^7^GpppN and m^7^GpppNm (N = C, U, G, A or m^6^A) from synthetic oligonucleotides, with the results showing quantitative release of all m^7^GpppN and m^7^GpppNm caps (**Supplementary Figure S10**). Finally, the stability of cap structures during NP1 digestion was verified by spiking cap standards into the RNA digestion reactions with subsequent HPLC purification and isotope-dilution LC-MS/MS analysis (**Figure 1D and E**).

Recently a new type of mRNA cap has been proposed containing m^1^A (36–38). These caps, m^7^Gpppm^1^A or m^7^Gpppm^1^Am, were predicted based on the binding of m^1^A antibodies to 5’ ends of mRNA (39). However, no biochemical validation was presented. To quantify these caps biochemically, we first wanted to develop cap purification protocols that would preserve m^1^A, due to the potential for this nucleotide to convert to m^6^A by the Dimroth rearrangement (**Supplementary Figure S7a**) (36–38), we defined the fate of m^1^A and m^1^Am ribonucleosides during the RNA isolation and processing. As shown in **Supplementary Figure S7b**, conversion of m^1^A to m^6^A occurred at each step – TRIzol RNA extraction (7%), polyA-tailed RNA purification (17%), GeneRead rRNA depletion (36%), and RNeasy MinElute Cleanup (72%). This means that for yeast and mouse RNA, which were processed with all steps, 86% of initial m^1^A would have been converted to m^6^A. With LODs of 0.68 fmol for m^7^Gpppm^1^A and 0.11 fmol for m^7^Gpppm^1^Am (**Supplementary Table S1**), m^1^A- and m^1^Am-containing caps present at 10 fmol per μg of RNA, which is the lowest level among all of the canonical caps in humans, mice, and yeast as discussed shortly, would remain detectable even with 90% loss. For human RNA, which was processed without rRNA depletion and the RNA cleanup steps, m^1^A and m^1^Am losses were at most 23%, so m^7^Gpppm^1^A and m^7^Gpppm^1^Am should be readily detectable in human mRNA if present.

Based on our validation steps, CapQuant was now applied to viral, bacterial, yeast, mouse, and human RNA to discover new cap structures, quantify m^1^A or m^1^Am in caps, and to define the composition and dynamics of the cap epitranscriptome.

### Quantitative analysis of the cap landscape in eukaryotic, prokaryotic, and viral RNA

With an optimized CapQuant method in hand, we applied it to define the landscape of caps in coding and non-coding RNAs from a range of organisms, including humans, mice, yeast, bacteria, and an RNA virus. Focusing first on poly(A)-tailed RNAs (mainly mRNA) from log-growing human CCRF-SB lymphoblasts (**Figure 2A**), we were able to quantify the components of the cap epitranscriptome. Of the 26 targeted caps, 10 were reproducibly detected for a total of 2,078 fmol of caps per μg of RNA. As expected, the five cap 1 structures (m^7^GpppNm) comprised the majority of all caps (88%, 1,830 fmol/μg RNA) with no cap 0 structures (m^7^GpppN) detected. Consistent with the fact that very few transcriptional start sites (TSS) in humans start with a uridine (**Figure 3A** and **Supplementary Figure S6b**), m^7^GpppUm comprised only 1% of second-nucleotide subtypes (**Figure 2A**), which ranged from 23 to 595 fmol/μg RNA. The most abundant caps were the C, G and A subtypes, found in nearly equal proportions: 33% m^7^GpppCm, 32% m^7^GpppGm, and 19% m^7^Gpppm^6^Am/15% m^7^GpppAm. This distribution correlates strongly with the distribution of predicted TSS (+1 position) frequencies in humans (**Figure 3A** and **Supplementary Figure S6b**). Our analysis further revealed four previously undescribed cap structures (**Figure 1A**): m^7^Gpppm^6^A, FAD, UDP-Glc, and UDP-GlcNAc. The m^7^Gpppm^6^A structure proved to be relatively abundant at 12% of all mRNA caps (244 fmol/μg RNA), which contradicts previous claims of the absence of this cap based on crude thin-layer chromatography analyses (14) and in a non-quantitative LC-MS assay (18). Additionally, this cap demonstrates that 2’-*O*-methylation is not essential in mRNAs, as has been previously suggested to suppress innate host antiviral responses (3). The structures of the four metabolite caps (NAD, FAD, UDP-Glc and UDP-GlcNAc) were unequivocally confirmed by three signature MRM transitions defined with standards (**Figure 1E** and **Supplementary Figure S8**). Compared to cap 1 structures, however, the levels of these metabolite caps were ~100-fold lower at 0.40-2.9 fmol/μg RNA (**Figure 2A** and **Table 1**). UDP-GlcNAc and NAD being the two most abundant structures is consistent with the relative abundance of these metabolites in human cells (40,41) and thus with the idea that nucleotide metabolites can initiate transcription (9). Notably, we were unable to detect m^7^Gpppm^1^A or m^7^Gpppm^1^Am in human mRNAs (**Supplementary Figure S11**).

**Figure 2.**
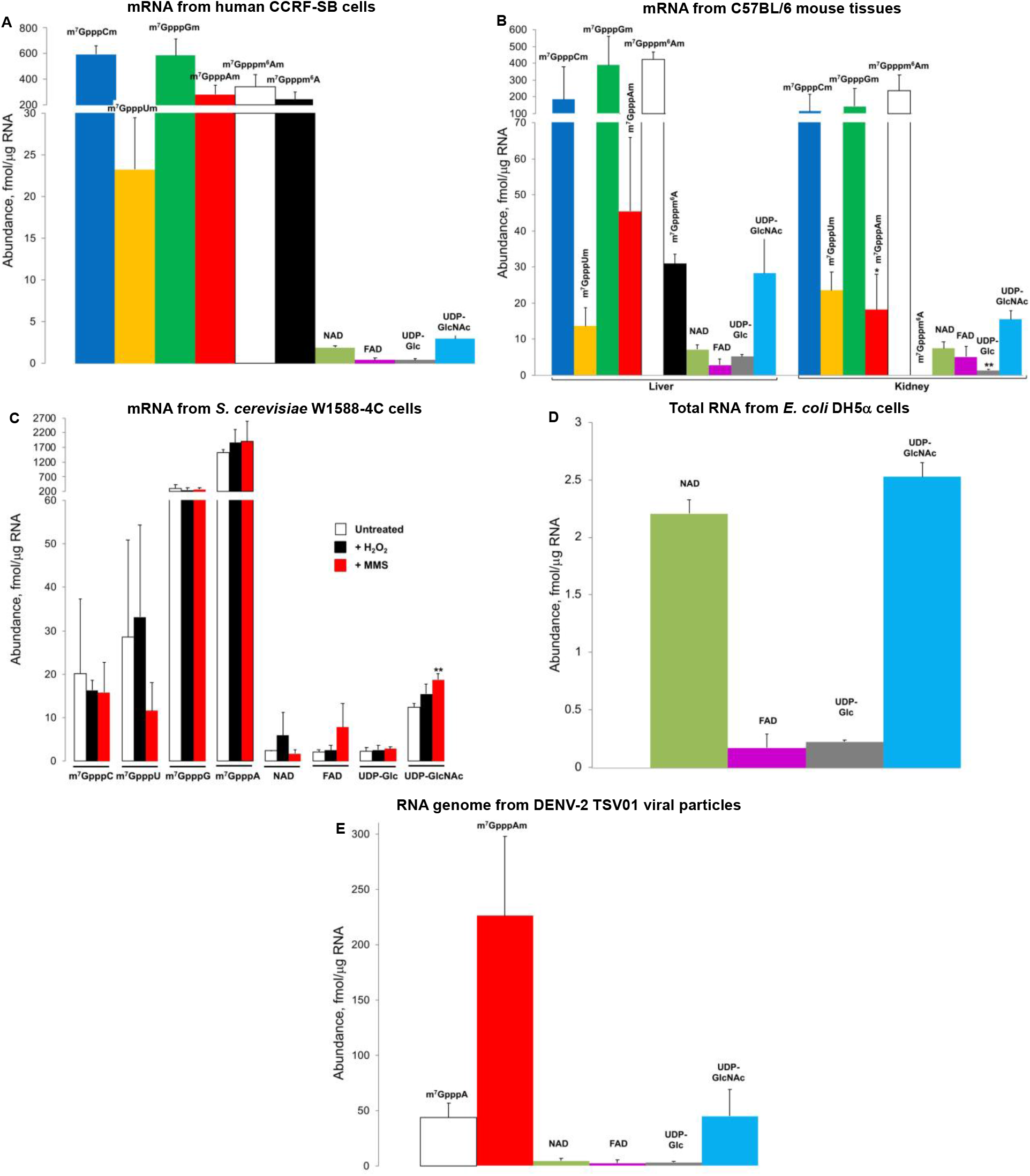
Quantification of 5’ cap structures in cellular RNA and viral RNA genome by CapQuant. (**A**) mRNA from Human CCRF-SB cells. (**B**) mRNA from mouse C57BL/6 liver and kidney tissues. * *P* < 0.05, ** *P* < 0.01, two-tailed paired Student’s *t* test. (**C**) mRNA from *Saccharomyces cerevisiae* W1588-4C cells. Exposure to hydrogen peroxide (H_2_O_2_) or methyl methanesulfonate (MMS) induces changes to the profile of 5’ cap structures in mRNA from *Saccharomyces cerevisiae*. From left to right: untreated, H_2_O_2_-treated, MMS-treated. ** *P* < 0.01, two-tailed unpaired Student’s *t* test. (**D**) *E. coli* DH5α total RNA. (**E**) DENV-2 virus RNA genome. Values represent mean ± SD for three independent cultures for CCRF-SB, W1588-4C and DH5α, for three biological replicates of three mice and H_2_O_2_- or MMS-treated W1588-4C cells, and for three technical replicates of a single culture for DENV-2.

**Figure 3.**
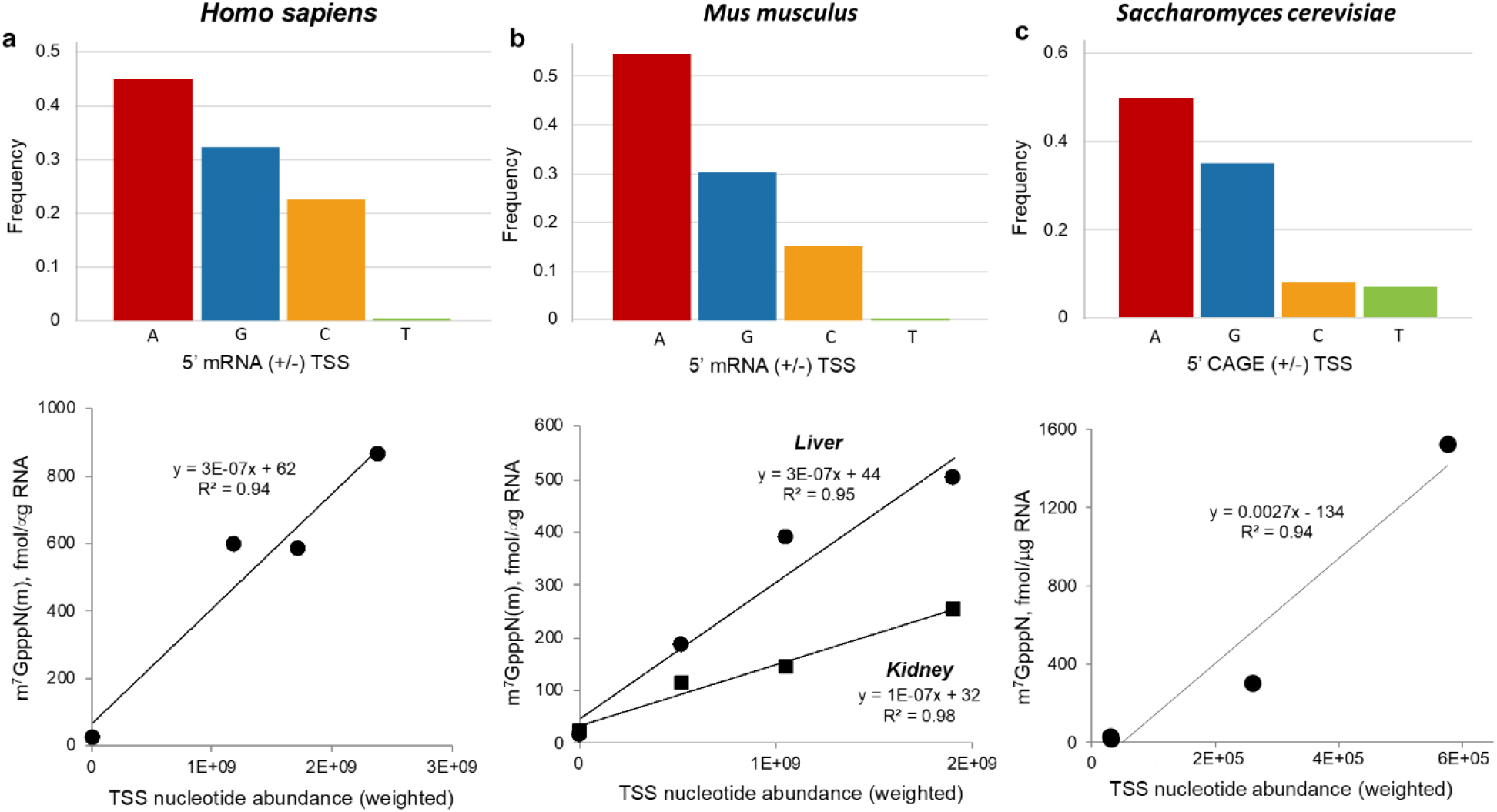
Cap profile correlation with CAGE-analyzed transcription start site (TSS) nucleotide distribution. The frequency of A, G, C, and T as the second nucleotide in m^7^GpppN caps was plotted against the distribution of these nucleotides at TSSs in (**A**) human (FANTOM5-weighted TSS), (**B**) mouse liver and kidney (FANTOM5-weighted TSS), and (**C**) *Saccharomyces cerevisiae* (YeasTSS-weighted TSS). TSS values were calculated as described in MATERIALS AND METHODS.

**Table 1:** Cap compositions in cellular and viral RNA species^1^

| Cap | Level, fmol per μg RNA<br>(Percentage, %) | | | | | |
| --- | --- | --- | --- | --- | --- | --- |
| | Human<br>CCRF-SB<br>mRNA | Mouse<br>C57BL/6<br>liver mRNA | Mouse<br>C57BL/6<br>kidney mRNA | *S. cerevisiae*<br>W1588-4C<br>mRNA | *E. coli*<br>DH5α<br>total RNA | DENV-2<br>TSV01<br>RNA genome |
| m<sup>7</sup>GpppCm | 595 ± 65<br>(27 ± 3) | 184 ± 195<br>(16 ± 17) | 114 ± 101<br>(20 ± 18) | nd | nd | nd |
| m<sup>7</sup>GpppUm | 23 ± 6<br>(1.1 ± 0.3) | 14 ± 5<br>(1.2 ± 0.5) | 24 ± 5<br>(4.2 ± 0.9) | nd | nd | nd |
| m<sup>7</sup>GpppGm | 585 ± 128<br>(28 ± 6) | 389 ± 172<br>(34 ± 15) | 144 ± 105<br>(25 ± 18) | nd | nd | nd |
| m<sup>7</sup>GpppAm | 282 ± 76<br>(14 ± 4) | 46 ± 20<br>(4.0 ± 1.8) | 18 ± 10<br>(3 ± 2) | nd | nd | 226 ± 72<br>(70 ± 22) |
| m<sup>7</sup>Gpppm<sup>6</sup>Am | 345 ± 93<br>(17 ± 4) | 425 ± 43.4<br>(38 ± 4) | 237 ± 92<br>(42 ± 16) | nd | nd | nd |
| m<sup>7</sup>GpppC | nd | nd | nd | 20 ± 17<br>(1.1 ± 0.9) | nd | nd |
| m<sup>7</sup>GpppU | nd | nd | nd | 28 ± 22<br>(1.5 ± 1.2) | nd | nd |
| m<sup>7</sup>GpppG | nd | nd | nd | 305 ± 130<br>(16 ± 7) | nd | nd |
| m<sup>7</sup>GpppA | nd | nd | nd | 1524 ± 106<br>(80 ± 6) | nd | 44 ± 12<br>(14 ± 4) |
| m<sup>7</sup>Gpppm<sup>6</sup>A | 244 ± 61<br>(12 ± 3) | 31 ± 3<br>(2.7 ± 0.2) | nd | nd | nd | nd |
| NAD | 1.9 ± 0.2<br>(0.09 ± 0.01) | 7.1 ± 1.2<br>(0.6 ± 0.1) | 7.4 ± 1.8<br>(1.3 ± 0.3) | 2.4 ± 0.1<br>(0.13 ± 0.01) | 2.2 ± 0.1<br>(43 ± 2) | 4.5 ± 2.5<br>(1.4 ± 0.8) |
| FAD | 0.4 ± 0.2<br>(0.02 ± 0.01) | 2.8 ± 1.8<br>(0.2 ± 0.2) | 5.0 ± 3.1<br>(0.9 ± 0.5) | 2.0 ± 0.6<br>(0.11 ± 0.03) | 0.17 ± 0.12<br>(3.3 ± 2.4) | 2.5 ± 2.6<br>(0.8 ± 0.8) |
| UDP-Glc | 0.4 ± 0.1<br>(0.02 ± 0.01) | 5.2 ± 0.5<br>(0.4 ± 0.04) | 1.4 ± 0.3<br>(0.25 ± 0.05) | 2.2 ± 0.8<br>(0.12 ± 0.04) | 0.22 ± 0.02<br>(4.3 ± 0.3) | 3.2 ± 0.6<br>(1 ± 0.2) |
| UDP-GlcNAc | 2.9 ± 0.4<br>(0.14 ± 0.02) | 28 ± 10<br>(2.5 ± 0.9) | 15 ± 2<br>(2.7 ± 0.4) | 12 ± 0.8<br>(0.66 ± 0.04) | 2.5 ± 0.1<br>(49 ± 2) | 44 ± 24<br>(14 ± 7) |
| Total caps,<br>fmol/μg RNA | 2078 ± 430 | 1131 ± 449 | 566 ± 320 | 1896 ± 278 | 5.1 ± 0.4 | 325 ± 114 |

<sup>1</sup> Values (as fmol per μg RNA or percentage for each detected cap) represent mean ± SD for three independent cultures for all cell lines, for tissues from three different mice, and for three technical replicates of a single culture for DENV-2 virus. nd, not detectable.

We next sought to understand whether the cap epitranscriptome is different in different cell types. The same 10 mRNA caps observed in the human cells were also found in mouse liver and kidney tissue mRNAs at 1,131 and 566 fmol/μg RNA, respectively. Mice similarly showed relatively low abundance of m^7^GpppUm and high levels of m^7^GpppGm and m^7^GpppCm (**Figure 2B** and **Table 1**), though m^7^GpppAm was >5-fold lower in mice liver and kidney than in human CCRF-SB cells (**Figure 2B** and **Table 1**). The large differences between the ratio of m^7^GpppAm and m^7^Gpppm^6^Am in different cell types supports a role for m^6^Am as a regulatable modification in mRNA. A comparison of caps in liver and kidney showed several striking tissue-specific differences, most notably the absence of detectable m^7^Gpppm^6^A in kidney (**Figure 2B** and **Table 1**). Other tissue-specific differences include >2-fold lower levels of m^7^GpppGm (p > 0.05), m^7^GpppAm (p > 0.05), m^7^Gpppm^6^Am (p < 0.05), and UDP-Glc (p < 0.01) in kidney compared to liver, and small variations in the levels of m^7^GpppCm, m^7^GpppUm, NAD, FAD and UDP-GlcNAc (**Figure 2B**). Similar to humans, the cap second nucleotide distribution correlates strongly with the distribution of predicted TSS frequencies in mice (**Figure 3B** and **Supplementary Figure S6c**).

In contrast to the cap 1 structures in mammalian cells, the only canonical caps in *S. cerevisiae* were the expected cap 0 structures (m^7^GpppN), with abundances between 20 and 1,524 fmol/μg RNA (**Figure 2C** and **Table 1**). m^7^GpppA constituted 80% of all caps (1,896 fmol/μg), with m^7^GpppA > m^7^GpppG (16%) >> m^7^GpppU (1.5%) > m^7^GpppC (1.1%). This distribution correlates strongly with the distribution of predicted TSS frequencies in *S. cerevisiae* (**Figure 3C** and **Supplementary Figure S6d**). The four nucleotide metabolite caps were present in the *S. cerevisiae* mRNAs at abundances from 2.0 to 12.4 fmol/μg RNA, which is higher than in humans and mice (**Figure 2A-B** and **Table 1**). Notably, we found no evidence for the presence of methylated forms of A in any cap structures in yeast.

The tissue-specific variations in cap structure and quantity in mice raised the possibility that cap landscape would vary as a result of stress-specific changes in gene expression. To this idea, we quantified the cap profile in yeast exposed to well-characterized oxidative and alkylation stresses caused by hydrogen peroxide (H_2_O_2_) and methyl methanesulfonate (MMS), respectively. Both treatments resulted in modest changes in the levels of several caps (**Figure 2C**), with a significant increase in the level of UDP-GlcNAc cap (p < 0.01). However, there were no striking changes in cap levels for these two stressors.

As expected, the m^7^G-type cap structures typical of eukaryotes were not detectable in the total RNA from *E. coli* (**Figure 2D** and **Table 1**). Here we analyzed total RNA instead of mRNA because of the low prevalence of polyA tails in the *E. coli* mRNA pool, with only 2-50% of mRNAs shown to have polyA that are generally short at 14-60 nt (42). While NAD and UDP-GlcNAc were the major metabolite caps, which is consistent with the relatively high concentration of these metabolites in *E. coli* (43), the four metabolite caps in *E. coli* occurred at 10-fold lower levels than in yeast, ranging from 0.20 to 2.5 fmol/μg RNA (**Figure 2D** and **Table 1**). This suggests differing propensities of the yeast and bacterial RNA polymerases for using nucleotide metabolites to initiate transcription.

Finally, in dengue purified virion RNA genomes, the total level of detected caps amounted to 325 ± 114 fmol/μg RNA. This is consistent with nearly all copies of the ~10,700 nt RNA genome (288 fmol/μg RNA) possessing a cap. The major cap structure (70%) was found to be the cap 1 m^7^GpppAm at 226 fmol/μg RNA (**Figure 2E** and **Table 1**). Surprisingly, the cap 0 structure m^7^GpppA represented 14% of all caps. The abundance of the four metabolite caps ranged from 2.5 to 44.8 fmol/μg RNA, which is similar to yeast.

## DISCUSSION

Here we present CapQuant, an analytical method combining off-line HPLC enrichment with isotope-dilution LC-MS/MS analysis for analysis of the diversity and dynamics of the cap epitranscriptome. This method overcomes the shortcomings of existing cap analysis tools, which are limited to individual cap structures (20,44,45), are poorly quantitative (5,14,18), and lack of chemical specificity (5,14), to enable accurate, specific and sensitive quantification of the RNA cap landscape in any organism. It achieves high-coverage with absolute quantification – a key feature of the method – over a broad dynamic range starting at attomole levels (as little as 600 ng of RNA) and the capacity to expand to other new RNA cap structures, including the methylated guanosine caps observed in pre-tRNA (21). While isotope-labeled internal standards provide highly accurate absolute quantification, rigorous cap quantification can still be performed with external calibration curves using unlabeled standards or even with other chemically similar cap standards. The use of off-line ion-pairing HPLC (46) for cap enrichment (**Figure 1B**) greatly enhances quantitative sensitivity by reducing interference from the matrix and non-cap nucleotides. It further helps in new cap discoveries akin to DNA “adductomics” (47) by collecting ion-pairing HPLC fractions across the elution time-course and analyzing them by MS scanning for novel MS signals. However, as the use of ion-pairing agents involves chronic contamination of HPLC and MS systems, a dedicated HPLC system and volatile ion-pairing agents for its complete removal before LC-MS/MS analysis is recommended.

Application of CapQuant to eukaryotic and flavivirus RNA has demonstrated that the composition of RNA caps varies between different tissues, supporting the idea of a regulated cap epitranscriptome. In addition, our data (i) quantitatively confirmed previous qualitative observations about the predominance of m^7^G-type caps, (ii) confirmed the lack of GpppN caps, and (iii) facilitated the discovery of novel and noncanonical caps, such as the metabolite caps (10,12,48), (iv) suggest that cap m^1^A//m^1^Am is unlikely to be present at appreciable levels in mRNA caps, raising a warning on interpreting any data on m^1^A//m^1^Am in RNA, (v) revealed the occurrence of surprisingly high proportions of caps lacking 2’-*O*-methylation in mammalian mRNA and viral RNA genome, and (vi) facilitated transcription start site analysis.

The lack of detectable GpppN caps could reflect the cap quality control system described in mammalian (DXO/Dom3Z protein) and yeast cells (Rai1–Rat1 and Dxo1). These systems possess decapping, pyrophosphohydrolase, and 5’-to-3’ exonuclease activities that appear to target caps lacking m^7^G (49,50).

With regard to m^1^A, two antibody-based methods concluded that m^1^A was widespread in mammalian mRNA (36,37), with subsequent studies proposing that m^1^A could exist as part of a novel cap structure comprising m^7^Gpppm^1^A or m^7^Gpppm^1^Am (39). However, biochemical studies were not used to demonstrate the existence of these novel mRNA caps. Another study used an antibody and sequencing-based approach to detect m^1^A-induced reverse transcriptase errors and suggested that m^1^A was present at much lower levels than previously thought (38). Here we demonstrate that m^1^A is unlikely to be present at appreciable levels in mRNA caps, as least in cultured human lymphoblasts. Even after quantitatively accounting for artifactual loss of m^1^A by Dimroth rearrangement to m^6^A and optimizing the CapQuant method to minimize this conversion, we did not detect any m^7^GpppN or m^7^GpppNm caps containing m^1^A or m^1^Am. Our data suggest that the levels of m^1^A and m^1^Am at mRNA caps, if they exist, are below the LODs which are 0.68 fmol and 0.11 fmol respectively. Thus, although this study does not rule out the existence of m^1^A//m^1^Am at mRNA caps, they are below the limits of detection, which suggests that they, if present, are found in less than 1/16,000 and 1/100,000 mRNA transcripts respectively from cultured human lymphoblasts. It should be noted that we did not attempt to solve the Dimroth rearrangement problem, thus we cannot be sure about the cap m^1^A//m^1^Am status. However, our observations raise a warning on interpreting any data on m^1^A//m^1^Am in RNA. While previous sequencing-based methods reported about a couple dozens of cap m^1^A in the HEK293T cells (38), the existence of cap m^1^A/m^1^Am in different tissues or samples needs further investigation.

In terms of novel cap discovery, we detected m^7^Gpppm^6^A as a novel cap in mRNA from human cells and mouse liver (**Figure 2A-B** and **Table 1**). The presence of m^7^Gpppm^6^A in mouse liver but not kidney points to a tissue-specific role for this cap. This could arise by demethylation of the m^7^Gpppm^6^Am cap through a yet unknown demethylase, or by *N*^6^-methylation of adenosine at the first transcribed nucleotide in mRNAs independent of the adenosine 2’-*O*-methylation status. Indeed, recent *in vitro* biochemical studies have shown that PCIF1, the enzyme responsible for synthesis of m^6^Am in mRNA caps, can also act on m^7^GpppA-capped mRNA to form m^7^Gpppm^6^A-capped mRNA (18,51,52). Thus, in cells, m^7^GpppA caps might undergo either 2’-*O*-methylation, *N*^6^-methylation, or both.

CapQuant also expanded the repertoire of 5’ cap structures with the discovery of three novel metabolite caps (FAD, UDP-Glc and UDP-GlcNAc) in all the RNA species analyzed (**Figures 1** and **2**). This expands the generality of the idea that nucleotide metabolites can serve as caps in cellular and viral RNA (2). However, metabolite caps (NAD, FAD, UDP-Glc and UDP-GlcNAc) are rare in eukaryotes, accounting for 0.3-5.1% in total of all caps detected (**Figure 2** and **Table 1**) across eukaryotic cells and tissues. There is a strong stochastic basis for metabolite caps formation due to (i) their low abundance relative to the NpppN canonical caps in eukaryotes (>10-fold lower; 0.2-20 fmol/μg *versus* 10-600 fmol/μg), (ii) the similar frequencies of each cap type in all organisms, (iii) the variation in metabolite cap levels among tissues and stresses, and (iv) their proportionality to cellular metabolite pools. The role of nutrient availability and metabolite pools as determinants of metabolite cap levels is illustrated by several studies. First, it was shown by Walters *et al*. (53) that there were more NAD-capped mRNAs in *S. cerevisiae* grown in minimal medium compared to rich YEPD medium, which suggests that the levels of NAD caps are sensitive to nutrient status. Similarly, Canelas *et al*. found that NAD levels in *S. cerevisiae* are sensitive to culturing conditions and nutrient status (54). This variability in metabolite levels as a determinant of metabolite cap levels may explain the 33-fold difference in NAD caps observed here and in the studies of Grudzien-Nagolska *et al*. in *S. cerevisiae* (45), though contributions from the different analytical methods could also account for the different NAD cap levels. Finally, Grudzien-Nagolska *et al*. demonstrated that changes in cellular NAD levels in HEK293T cells correlate with changes of the levels of NAD caps (45). These studies all show a variability in metabolite cap levels based on metabolite pool levels in a way that suggests a potential signaling or regulatory function of metabolite caps. An emerging literature supports this idea. For example, the NAD cap has been shown to be present on a subset of mRNAs that are targeted for rapid decay in mammalian cells (11,55), while Kiledjian and coworkers have observed a post-transcriptional NAD capping activity, which suggests that this cap is not simply a transcriptional mistake (11).

The potential for variation in metabolite cap levels as a function of cell state is also illustrated with viral infections. For example, human cytomegalovirus (HCMV) infection upregulates UDP-GlcNAc levels in host cells (56) with similar metabolic shifts observed in other viruses (57,58). Hence it is proposed that dengue infection upregulates host cellular UDP-GlcNAc levels, especially since viral envelope (E) protein *N*-glycosylation is partly derived from UDP-GlcNAc in host cells (59,60). Higher host cell levels of UDP-GlcNAc may lead to increased transcription initiation with this nucleotide metabolite, which would explain the relatively large proportion of UDP-GlcNAc-capped viral transcripts detected in dengue purified virions (**Figure 2E** and **Table 1**). While the biological function of these metabolite caps requires further examination, RNA Pols appear to be capable of initiating transcription with the four nucleotide metabolites studied here and that dengue virus NS5 polymerase could initiate transcription with the metabolite caps in the same manner as the host RNA Pol. However, the ability of the metabolite-capped viral genomes to sustain viral replication is unknown. In addition to the above question regarding biological function, the discovery of the three novel metabolite caps also raises several other important questions. For example, can metabolite caps be exported from the nucleus into the cytoplasm in eukaryotic cells? Are metabolite caps found in RNAs that associate with polysomes? We think that answers to these questions can be readily obtained by directly applying CapQuant to relevant systems, i.e., RNA preparations from the nucleus and cytoplasm from the same population of cells, and polysome-bound RNAs.

Consistent with published observations, we found that the cap on the dengue RNA genome isolated from purified virions contained Am but not m^6^Am (**Figure 2E** and **Table 1**) as compared with human mRNA (12,61). CapQuant revealed that >30% of the viral particles generated during an infection possess caps that are counterproductive for viral replication and survival in the host: presumably untranslatable metabolite caps or the m^7^GpppA cap that activates innate immunity (**Figure 2E**). With an estimated single copy of the RNA genome per viral particle (62) and one viral particle infecting a host cell, the varying viral cap structures detected suggest that infections will occur with viral genomes having different translational efficiency or propensity to activate the antiviral response pathways. The fate of these variously capped viral genomes in the host is largely known. Indeed, there is controversy concerning the presence of m^6^Am in the caps on dengue-derived mRNAs isolated from infected cells, which presumably arise by replication of the infective genomic RNA (12,61,63). The sole published experimental work showed that only Am is found in dengue mRNA caps (63). The variable detection of m^6^Am in dengue mRNA caps could be explained by contamination with the abundance of host mRNA containing m^6^Am (**Figure 2A** and **Table 1**) or by *N*^6^-methylation of viral genomes and/or mRNA by host enzymes PCIF1 (18,51,52). Our observation that dengue genomic RNA present in purified virions lacks m^6^Am in the cap implies that any *N*^6^-methylation of Am in caps, if required for translation, must occur in viral transcripts used for protein production. However, replicated RNA genomes destined for virion assemblies can only possess m^7^GpppAm, m^7^GpppA, and the metabolite caps, as we observed (**Figure 2E** and **Table 1**). While some studies suggest that cap m^6^Am stabilizes a subset of mRNAs (24), other studies did not observe this effect (18,64,65). Interestingly, *N*^6^-methylation of A within the viral mRNA has been found to negatively regulate viral infection by reducing viral particle formation (66), while we have previously demonstrated that Am is present throughout the RNA genome of purified dengue virions (23). Clearly, there is significant work to be done to clarify the capping mechanisms involved in the various steps of viral infection.

CapQuant also showed that 14% of dengue genomes possess m^7^GpppA cap, and that 12% of human and 3% of mouse liver mRNAs possess m^7^Gpppm^6^A caps (**Figure 2** and **Table 1**). Although the observation of the latter stands in contrast to the inability to detect it in a crude, chemically-non-specific TLC method (5) or in insensitive LC-MS studies lacking standards (18), the detection of m^7^Gpppm^6^A caps is rigorously established here based on chromatographic and structural identity with a synthetic standard. The presence of m^7^Gpppm^6^A caps in human and mouse liver mRNAs is unlikely due to inefficient cellular 2’-*O*-methyltransferase activities or insufficient cellular innate immunity targeting cap 0 structures (67) since none of the other cap 0 structures were detectable in these RNAs, even in human CCRF-SB mRNA where the levels of m^7^GpppCm, m^7^GpppGm and m^7^GpppAm were almost twice or comparable to the level of m^7^Gpppm^6^Am (**Figure 2** and **Table 1**). Thus, these data suggest that at least in some cell types 2’-*O*-methylation is not present in all mRNAs, potentially suggesting that there may be specific cellular contexts in which 2’-*O*-methylation is not needed to suppress the innate host antiviral response (3). It is well established that RIG-I and MDA5 are sensors of non-self RNA in mammalian cells, and the IFIT complex is a dual sensor-effector of a cellular innate defense system (67) for caps without 2’-*O*-methylation. IFIT complex recognizes m^7^GpppA cap structures to inhibit translation of the viral genome during viral infection (3) while RIG-I binds to dsRNA with 5’-ppp and cap 0 (68). 2’-*O*-methylated caps (5’-pppNm and m^7^GpppNm) significantly reduces RIG-I binding affinity to target RNA and the innate defense system activation (68). This raises an important question: Does the proportion of m^7^GpppA caps present during a dengue infection correlate with virulence? It is reasonable to hypothesize that the more virulent dengue strains have evolved to minimize the proportion of m^7^GpppA caps that activate the innate antiviral response in host cells, a hypothesis readily tested by application of CapQuant to clinical dengue isolates replicated in culture. It is worth noting that previous efforts using two-dimensional TLC- or LC-MS-based methods did not detect m^7^Gpppm^6^A cap in mRNA from human cells (5,18). Although it is possible that m^7^Gpppm^6^A cap was indeed absent in those RNA preparations, the failure to detect this cap could also be due to lack of chemical specificity and insufficient sensitivity of the two-dimensional TLC method (5) or due to lack of sensitivity and selected monitoring of m^7^G-capped dimers (m^7^GpppN_1_Gp) to pentamers (m^7^GpppN_1_N_2_N_3_N_4_Gp) containing 0-3 methyl groups, which include only a portion of all possible m^7^G-capped sequences with A or methylated A as the first transcribed nucleotide, in the LC-MS method (18). Interestingly, the level of m^7^Gpppm^6^A cap in mRNA differed significantly among human CCRF-SB, mouse liver and mouse kidney (**Figure 2A-B**). To explore if these differences is linked to possible differences in expression of relevant cap modification enzymes, we assessed the relative expression of a selection of RNA cap modification enzymes including PCIF1, FTO, DCP2 and CMTR1 as well as ALKBH5 in the total RNA from CCRF-SB cells and mouse liver and kidney tissues on the transcription level by RT-qPCR. We observed no statistically significant difference between any two of these three samples in the relative expression of any enzyme examined (**Supplementary Figure S12**), suggesting that the differences in the level of m^7^Gpppm^6^A cap are likely due to other factors. We speculate that two such factors are secondary structure of 5’ ends of mRNAs (69) and helicase activity that is critical for CMTR1-mediated 2’-*O*-methylation of cap 1 in mRNAs harboring highly structured 5’ ends (70). Further studies are needed to fully address this question.

CapQuant analysis also provided strong corroboration for TSS studies, which are challenging due to the lack of long and conserved consensus sequences for TSSs. m^7^G caps with a purine as the first transcribed nucleotide represented the major caps found in mRNAs from human CCRF-SB (70%), mouse liver (82%) and kidney (74%) tissues, and *S. cerevisiae* W1588-4C (97%), with the relative abundance of different m^7^GpppNm’s or m^7^GpppN’s varying across the organisms and tissues (**Figure 2A-C** and **Table 1**). This preference for purines at the penultimate position in m^7^GpppN caps is rationalized by the strong preference for pyrimidine-purine dinucleotides at −1 and +1 positions of TSSs in the coding strand of eukaryotes, bacteria and some viruses, which is argued to facilitate the loading of ATP or GTP during transcription initiation (72–74). A comparison of the distribution of the second nucleotide in m^7^GpppN/m^7^GpppNm caps revealed by CapQuant to the distribution of TSSs (+1 position) predicted using the cap analysis gene expression (CAGE) method (29,75) was conducted for cross-validation. The CAGE method is advantageous over other TSS analysis methods in that it only captures capped transcripts and thus avoids false TSSs from degraded transcripts that do not contain caps. We observed a strong correlation between the cap second nucleotide distribution and the TSS distribution for *S. cerevisiae*, mice and humans (**Figure 3A-C** and **Supplementary Figure S6b-d**).

CapQuant is not without limitations. For example, the level of all caps per μg of mRNA in the mouse tissues was about 2- to 4-fold lower than in human cells and yeast. (**Table 1**). We cannot explain it, but could be due to presence of other types of caps not quantified in the present studies or a higher proportion of uncapped RNAs in the mouse tissue mRNA preparations. It is unlikely due to rRNA contamination as the Bioanalyzer profiles of all the cell and tissue mRNA preparations indicated undetectable level of rRNA contamination (**Supplementary Figure S4**). Also, it should be noted that in human, *S. cerevisiae* and *E. coli* cells, the levels of NAD cap revealed in the present study (**Table 1**) are up to 55-fold lower than those levels of the same cap determined or estimated in other studies (20,45). In addition to the variable accuracy of the different analytical methods, lower levels of NAD detection in the present studies could be due, at least in part, to differences in the cell culture conditions (45,53,54) and cell strains used in the different studies, as discussed earlier. In the case of *E. coli*, we analyzed caps in stationary-phase cells whereas Chen *et al*. analyzed caps in log-phase *E. coli* (20), which could contribute to the lower NAD cap level observed in our study. In addition, the non-significant changes in the abundance of NAD in the control experiments by Chen *et al*. (20) when spiking large amounts of NAD into the cell lysate prior to RNA isolation cannot rule out the possibility that the NAD they detected in the samples represented contaminating non-covalently bound NAD. CapQuant employs isotopically-labeled internal standards for cap quantification, which enhances the accuracy of the method.

In summary, beyond the applications in the quantification of cap structures in any type of RNA from *in viv*o or in vitro sources, CapQuant has wide potential use in many biological fields. Primarily, it can facilitate investigations into the dynamics, function and regulation of RNA caps in a broad wide range of biological processes and conditions. In addition, it can be readily applied to study RNA metabolism, such as RNA capping, RNA decapping and RNA decay. Notably, when combined with transcript-specific purification technology (76), it enables quantification of cap structures in specific transcripts and thus studies of transcript-specific capping and decapping, and gene-specific regulation. Finally, it permits investigations into the roles that cap-binding proteins, such as eIF4E and CBC, may play in the control of gene expression (77).

## SUPPLEMENTARY DATA

Supplementary Data are available online.

## ACKNOWLEDGEMENTS

We thank Profs. Graham Walker and Jianzhu Chen from the Massachusetts Institute of Technology for generously sharing *S. cerevisiae* W1588-4C and human CCRF-SB cells, respectively.

## FUNDING

This work was supported by grants from the National Institutes of Health (ES022858 to P.C.D. and CA186702 to S.R.J.), the Singapore-MIT Alliance for Research and Technology with a grant from the National Research Foundation of Singapore, the Inner Mongolia University with a grant from its “Steed Plan” High-Level Talents Program (awarded to J.W.), and the Nanyang Presidential Graduate Scholarship (B.L.A.C.).

## Conflict of interest statement

None declared.

