## Supplementary Materials for "Quantifying the RNA cap epitranscriptome reveals novel caps in cellular and viral RNA"

**Contents:**

Figures S1-S12
Tables S1-S3

- Figure S1: MS/MS spectra of cap nucleotides with assignments.
- Figure S2: LC-MS/MS analysis of RppH/SAP digestion mixtures.
- Figure S3: $^{1}$H NMR and $^{13}$C NMR spectra of the synthetic m$^{1}$Am.
- Figure S4: Bioanalyzer analysis of poly(A) RNA and DENV-2 viral RNA genome using the RNA 6000 Pico LabChips.
- Figure S5: Bioanalyzer analysis of total RNA using the RNA 6000 Nano or Pico LabChips.
- Figure S6: Transcription start site (TSS) nucleotide distribution in yeast, mice and humans.
- Figure S7: Dimroth rearrangement of m$^{1}$A and m$^{1}$Am during the RNA extraction, purification, cleanup and enzymatic digestion steps.
- Figure S8: Selective ion-chromatograms (SICs) for monitoring MRM transition(s) for cap structures.
- Figure S9: Calibration curves for the quantification of cap dinucleotides, m$^{1}$A, m$^{6}$A, m$^{1}$Am and m$^{6}$Am nucleosides.
- Figure S10: Quantification of the release of caps from m$^{7}$GpppN- and m$^{7}$GpppNm-capped RNA oligos during NP1 digestion (N = C, U, G, A or m$^{6}$A).
- Figure S11: Analysis of m$^{7}$Gpppm$^{1}$A and m$^{7}$Gpppm$^{1}$Am in RNA.
- Figure S12: Quantitative real-time PCR analysis of the expression of CMTR1, PCIF1, FTO, DCP2 and ALKBH5 genes in CCRF-SB, mouse liver, and mouse kidney total RNA.
- Table S1: Detection limits of known and potentially existing cap nucleotides:

- Table S2: Exact mass of synthetic unlabeled cap dinucleotides determined by high-resolution mass spectrometry.
- Table S3: Primers used for qPCR analysis.

**Figure S1.** MS/MS spectra of cap nucleotides with assignments. (**a−ab**) m$^7$GpppC, m$^7$GpppU, m$^7$GpppG, m$^7$GpppA, m$^7$Gpppm$^6$A, m$^7$GpppCm, m$^7$GpppUm, m$^7$GpppGm, m$^7$GpppAm, m$^7$Gpppm$^6$Am, GpppC, GpppU, GpppG, GpppA, Gpppm$^6$A, GpppCm, GpppUm, GpppGm, GpppAm, Gpppm$^6$Am, NAD, FAD, UDP-Glc, UDP-GlcNAc, m$^{2,2,7}$GpppG, dpCoA, m$^7$Gpppm$^1$A, m$^7$Gpppm$^1$Am. Ade, adenine; Gua, guanine; m$^7$Gua, *N*$^7$-methylguaninie; m$^6$Ade, *N*$^6$-methyladenine; m$^6$A, *N*$^6$-methyladenosine; Cyt, cytosine; G, guanosine; Flav, flavin; A, adenosine; N, nicotinamide; R, ribose; mR, 2-O-methylribose; H, hydrogen; p, phosphate; pp, diphosphate; ppp, triphosphate; N, nicotinamide; M, molecular ion; m$^1$Ade, *N*$^1$-methyladenosine; Ura, uracil; U, uridine; m$^{2,2,7}$Gua, *N*$^{2,2,7}$-trimethylguanine; m$^{2,2,7}$G, *N*$^{2,2,7}$-trimethylguanosine.

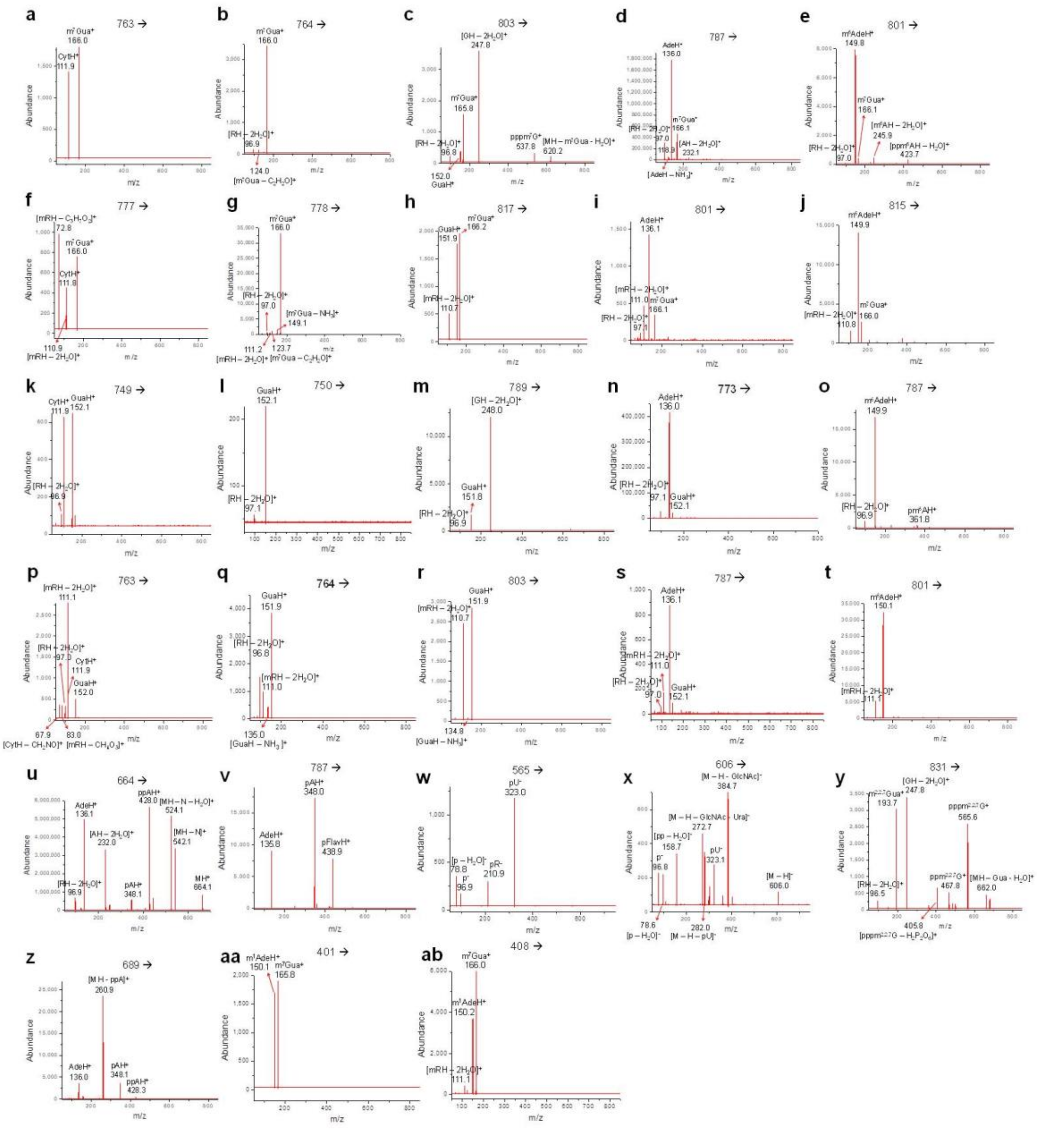

**Figure S2.** LC-MS/MS analysis of RppH/SAP digestion mixtures of (**a**) synthetic $m^7Gpppm^1A$ and (**b**) synthetic $m^7Gpppm^1Am$ and (**c**) a mixture of $m^1A$, $m^6A$, $m^1Am$ and $m^6Am$ standards.

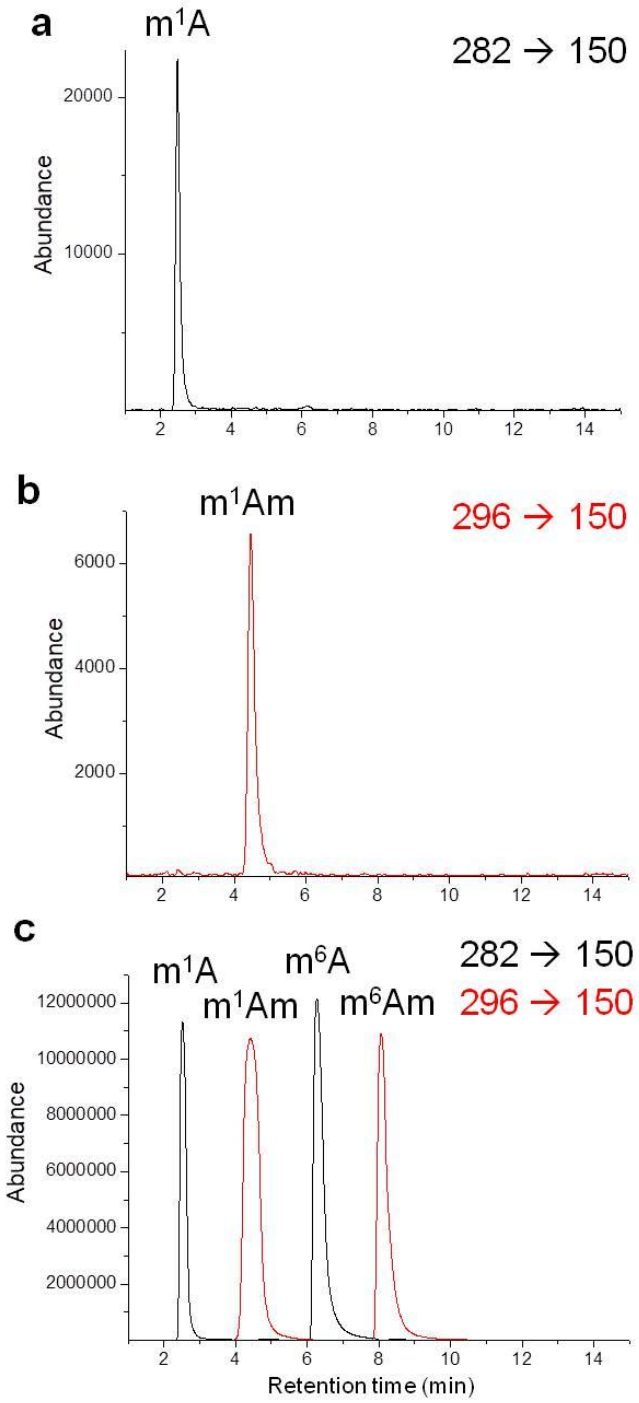

**Figure S3.** $^1$H NMR (**a**) and $^{13}$C NMR (**b**) spectra of the synthetic m$^1$Am.

**a**

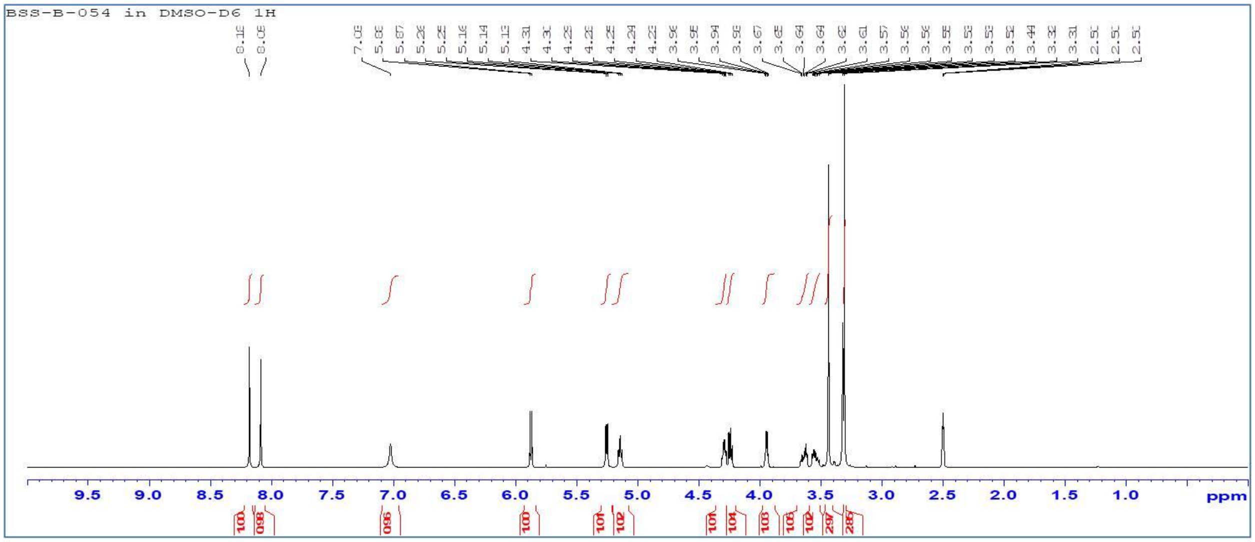


**b**

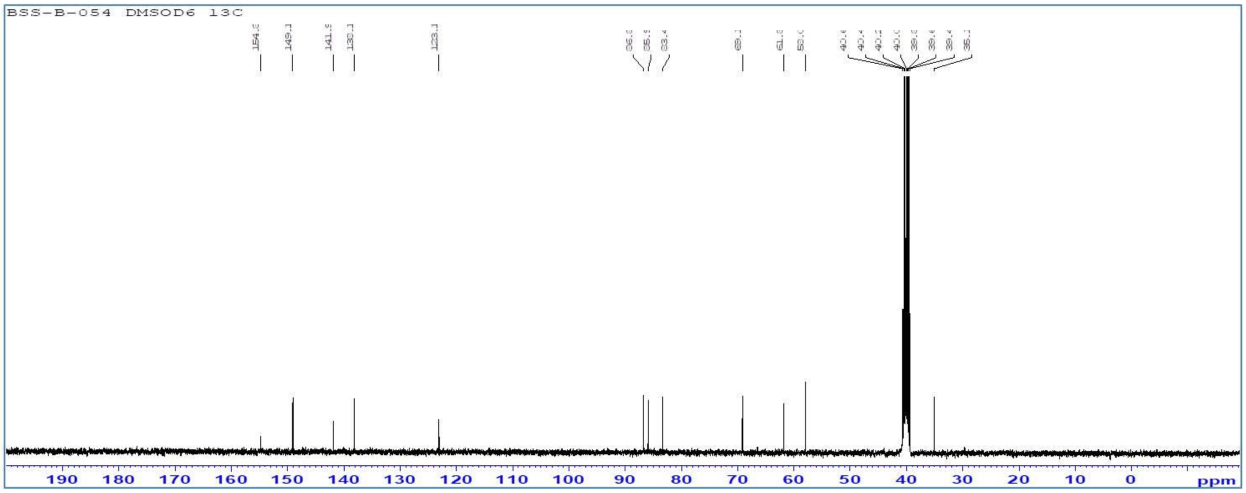

**Figure S4.** Bioanalyzer analysis of poly(A) RNA and DENV-2 viral RNA genome using the RNA 6000 Pico LabChips. (**a**) human CCRF-SB poly(A) RNA, (**b**) rRNA-depleted *S. cerevisiae* W1588-4C poly(A) RNA, (**c**) rRNA-depleted poly(A) RNA from $H_2O_2$-treated *S. cerevisiae* W1588-4C cells, (**d**) rRNA-depleted poly(A) RNA from MMS-treated *S. cerevisiae* W1588-4C cells, (**e**) rRNA-depleted poly(A) RNA from mouse C57BL/6 liver tissue, (**f**) rRNA-depleted poly(A) RNA from mouse C57BL/6 kidney tissue, (**g**) DENV-2 RNA genome before size-exclusion chromatography (SEC) purification, (**h**) DENV-2 RNA genome after SEC purification, (**i**) 6000 Pico RNA ladder.

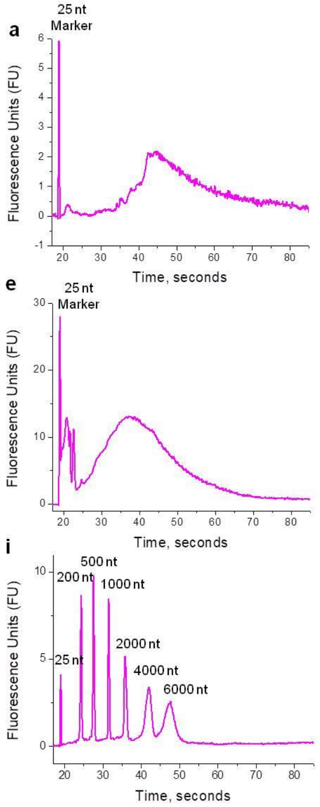

**Figure S5.** Bioanalyzer analysis of total RNA using the RNA 6000 Nano or Pico LabChips. (**a**) human CCRF-SB total RNA, (**b**) *S. cerevisiae* W1588-4C total RNA, (**c**) total RNA from $H_2O_2$-treated *S. cerevisiae* W1588-4C cells, (**d**) total RNA from MMS-treated *S. cerevisiae* W1588-4C cells, (**e**) *E. coli* DH5α total RNA, (**f**) mouse C57BL/6 liver tissue total RNA, (**g**) mouse C57BL/6 kidney tissue total RNA, and (**h**) 6000 Nano RNA ladder.

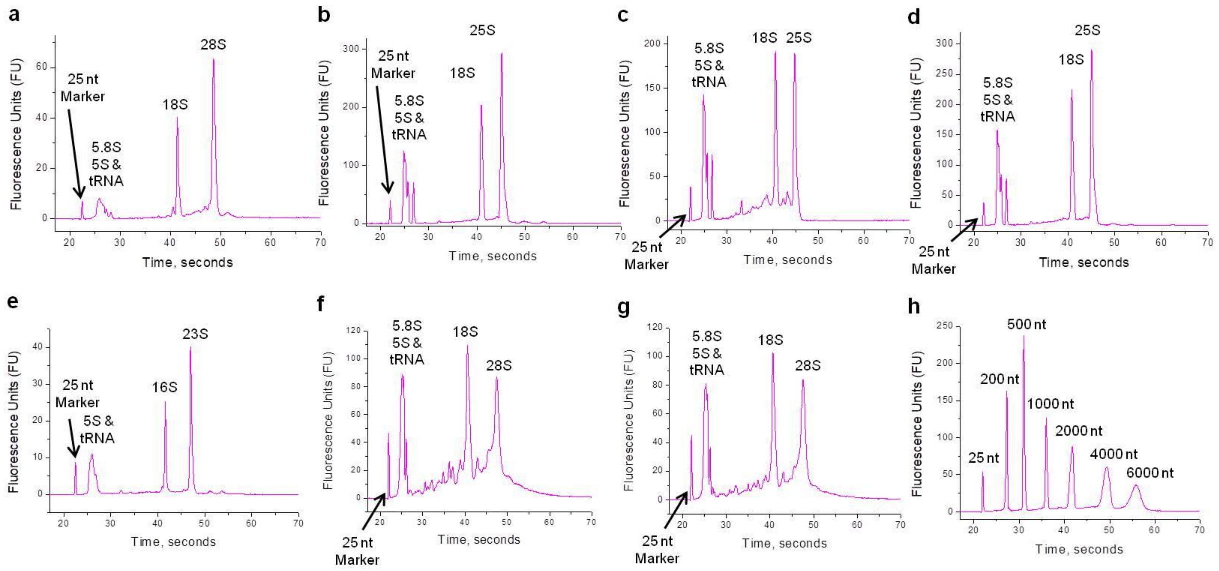

**Figure S6.** Transcription start site (TSS) nucleotide distribution in yeast, mice and humans. (**a**) workflow outline, (**b**) FANTOM5 5' CAGE TSS nucleotide distribution in humans, (**c**) FANTOM5 5' CAGE TSS nucleotide distribution in mice, (**d**) YeasTSS atlas 5' CAGE TSS nucleotide distribution.

a

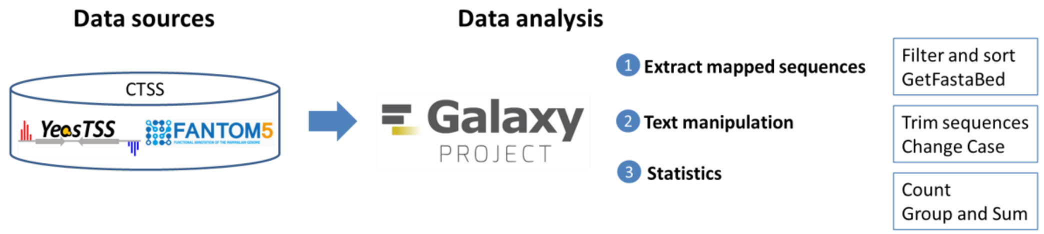


b

| hg38 FANTOM5 CAGE TSS (+/-) unweighted | | |
|---|---|---|
| A | 110766 | 52.77% |
| G | 61419 | 29.26% |
| C | 36091 | 17.19% |
| T | 1635 | 0.78% |
| Total | 209911 | |

| hg38 FANTOM5 CAGE TSS (+/-) weighted | | |
|---|---|---|
| A | 2386361482 | 44.94% |
| G | 1715089094 | 32.30% |
| C | 1189692638 | 22.41% |
| T | 18432901 | 0.35% |
| Total | 5309576115 | |

c

| mm10 FANTOM5 CAGE TSS (+/-) unweighted | | |
|---|---|---|
| A | 87031 | 52.85% |
| G | 46070 | 27.98% |
| C | 29856 | 18.13% |
| T | 1715 | 1.04% |
| Total | 164672 | |

| mm10 FANTOM5 CAGE TSS (+/-) weighted | | |
|---|---|---|
| A | 1906311037 | 54.45% |
| G | 1056943034 | 30.19% |
| C | 527507968 | 15.07% |
| T | 10510323 | 0.30% |
| Total | 3501272362 | |

d

| SacCer3 YeasTSS CAGE TSS (+/-) unweighted | | |
|---|---|---|
| A | 31287 | 49.81% |
| G | 22020 | 35.06% |
| C | 5061 | 8.05% |
| T | 4446 | 7.08% |
| Total | 62814 | |

| SacCer3 YeasTSS CAGE TSS (+/-) weighted | | |
|---|---|---|
| A | 576269.1867 | 64.16% |
| G | 259907.1422 | 28.94% |
| C | 31920.02775 | 3.55% |
| T | 30063.89398 | 3.35% |
| Total | 898160.2507 | |

**Figure S7.** Dimroth rearrangement of m$^1$A and m$^1$Am during the RNA extraction, purification, cleanup and enzymatic digestion steps. (**a**) Schematic illustration of m$^1$A-to-m$^6$A conversion (Dimroth rearrangement) under alkaline conditions at elevated temperatures. (**b**) Percentages of the Dimroth rearrangement of m$^1$A and m$^1$Am during each step. (**c**) LC-MS/MS analysis of m$^1$A, m$^6$A, m$^1$Am and m$^6$Am.

**a**

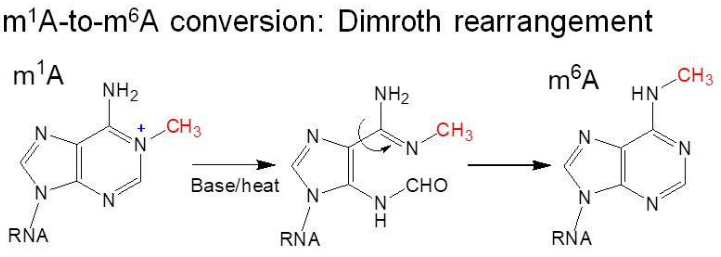


**b**

| Experimental steps | Dimroth rearrangement rate, % | |
|---|---|---|
| | m$^1$A → m$^6$A | m$^1$Am → m$^6$Am |
| TRIzol RNA extraction | 7.1 | 0.0 |
| MAG MAXI mRNA isolation | 17 | 2.0 |
| Generead rRNA depletion | 36 | 39 |
| RNeasy MinELute Cleanup | 72 | 50 |
| NP1 digestion | 21 | 2.7 |

**c**

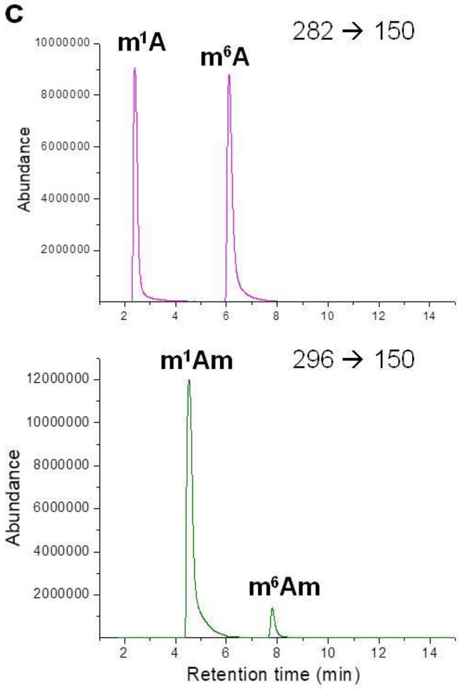

**Figure S8.** Selective ion-chromatograms (SICs) for monitoring MRM transition(s) for cap structures. (**a**-**x**) m$^7$Gpppm$^6$A, m$^7$GpppC, m$^7$GpppU, m$^7$GpppG, m$^7$GpppA, m$^7$GpppCm, m$^7$GpppUm, m$^7$GpppGm, m$^7$Gpppm$^6$Am, GpppC*, GpppU*, GpppG*, GpppA*, Gpppm$^6$A*, GpppCm*, GpppUm*, GpppGm*, GpppAm*, Gpppm$^6$Am*, dpCoA*, m$^{2,2,7}$GpppG*, FAD, UDP-Glc, and UDP-GlcNAc. For each cap, there are three panels: top, unlabeled standard; middle, analyte in RNA sample; and bottom, isotope-labeled standard spiked into RNA sample. Red asterisks denote caps that were not detectable in any of the RNA samples analyzed in this work.

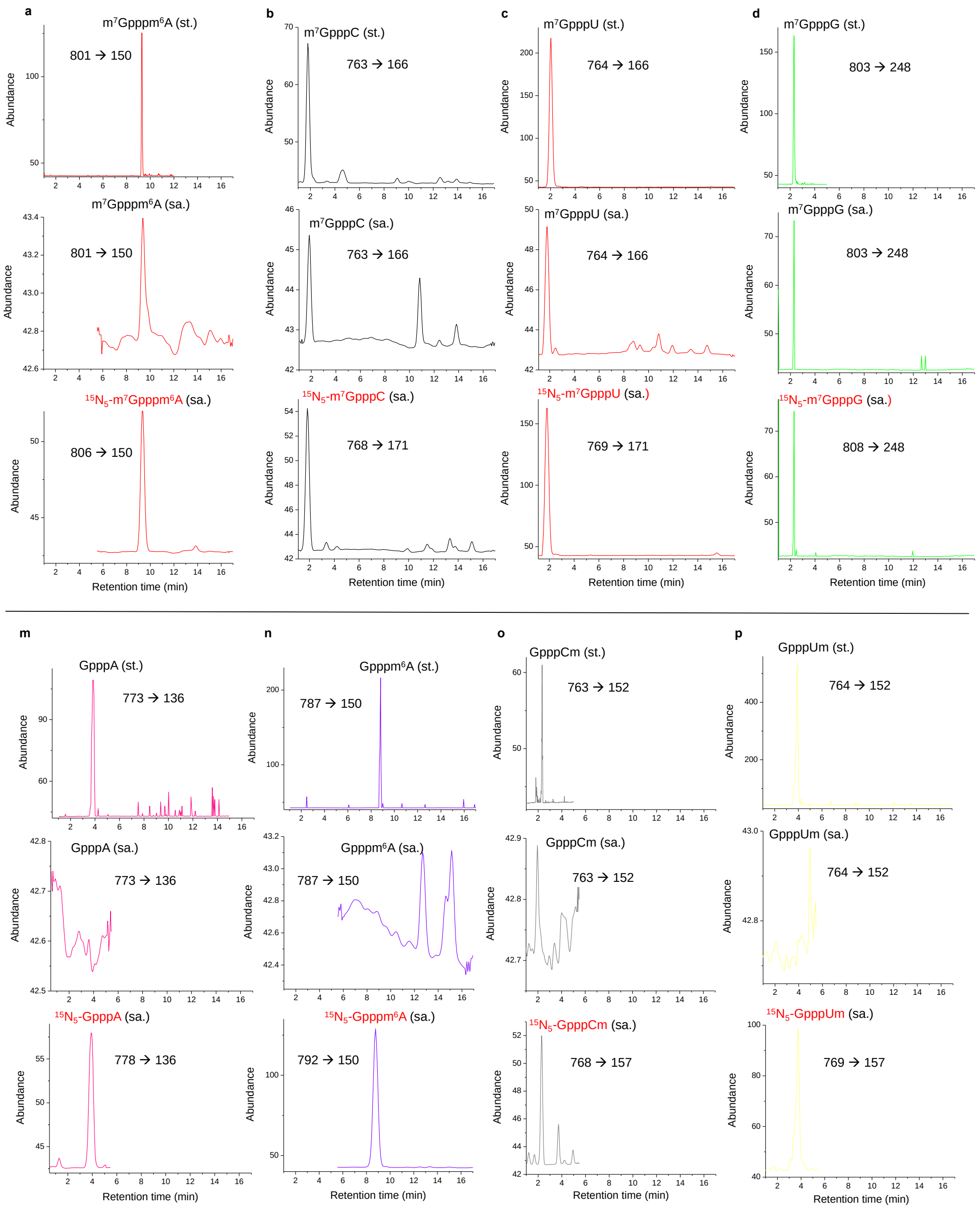

### Figure S8, continued

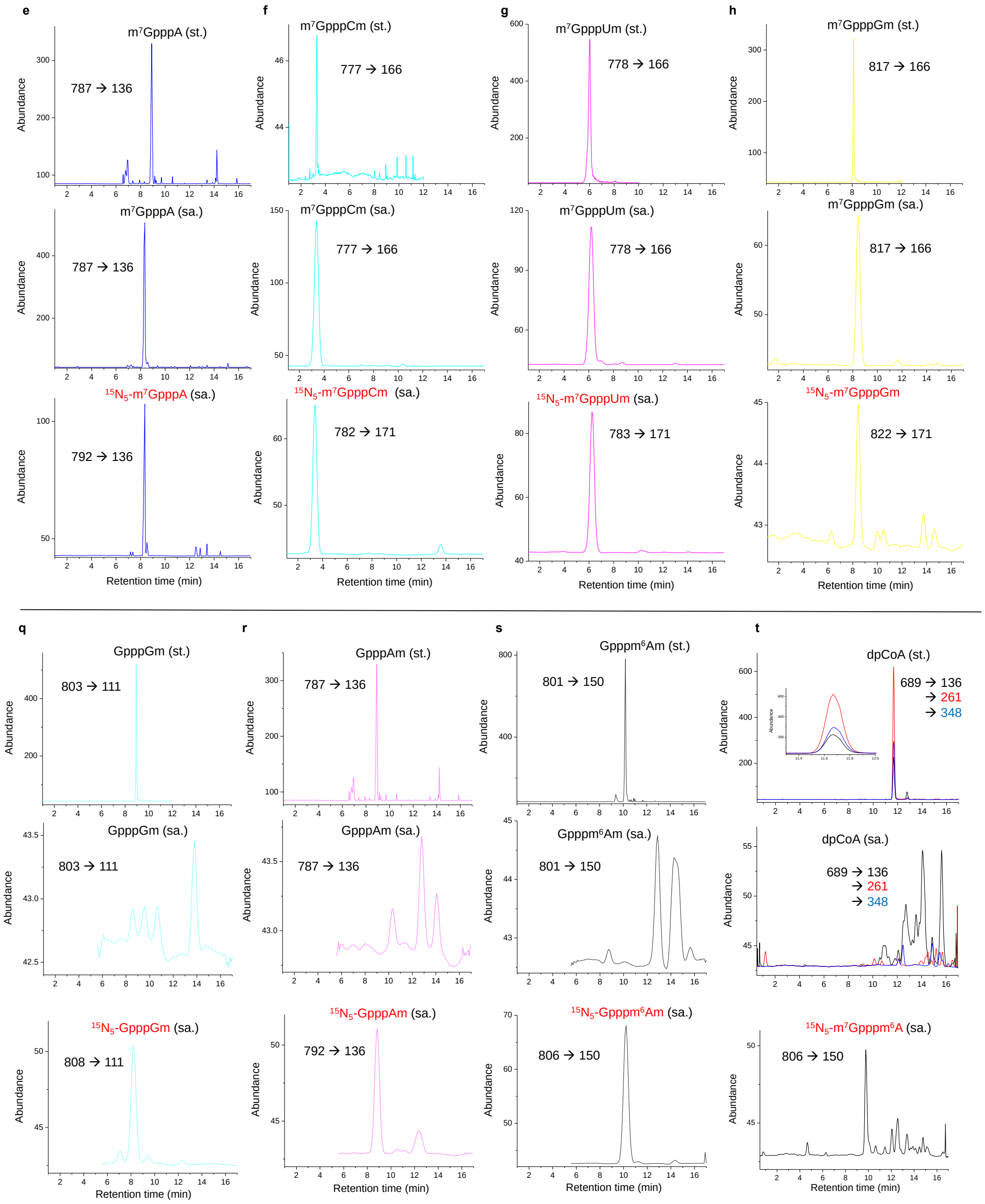

### Figure S8, continued

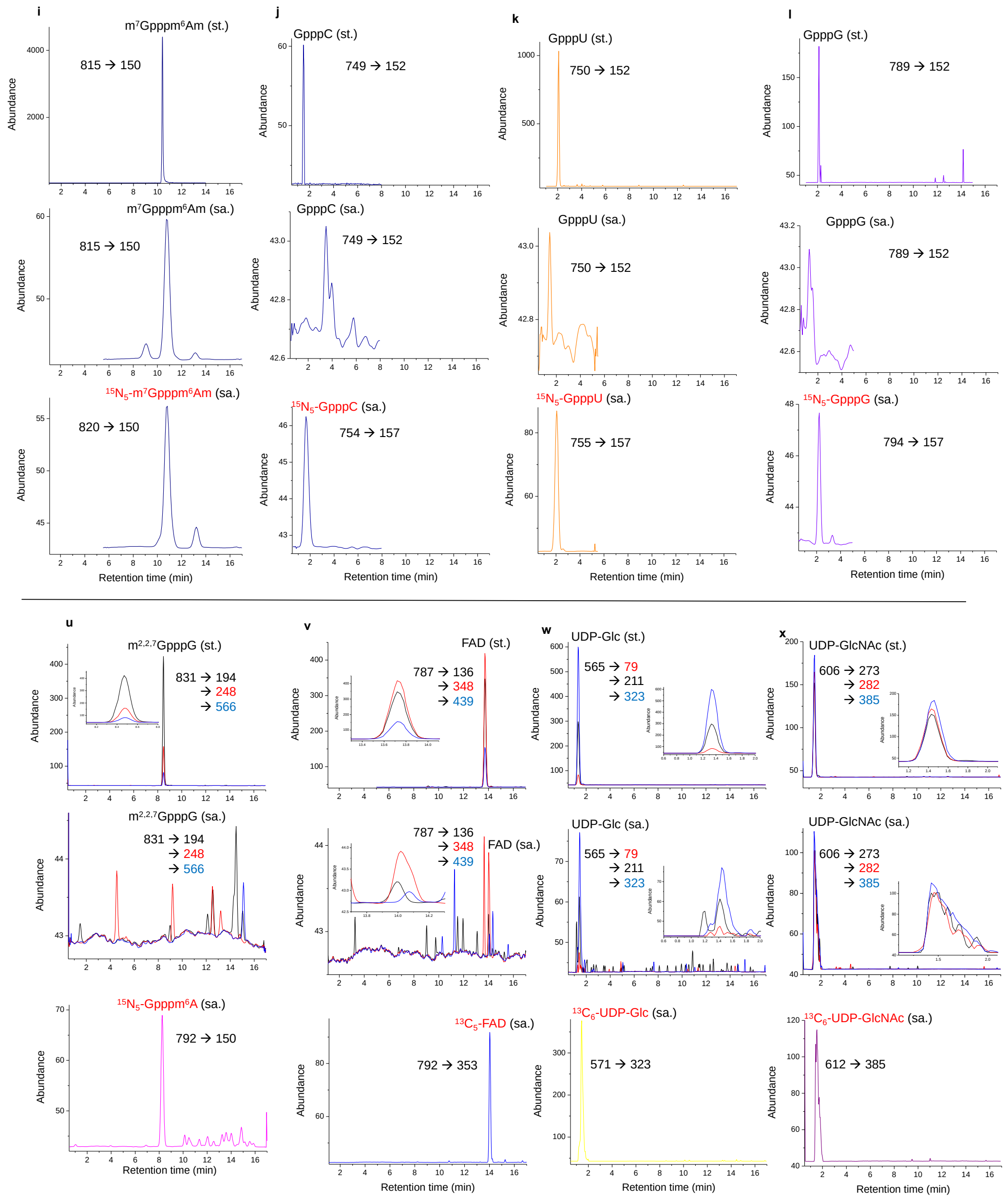

**Figure S9.** Calibration curves for the quantification of cap dinucleotides, m$^1$A, m$^6$A, m$^1$Am and m$^6$Am nucleosides. (**a**-**r**) m$^7$GpppC, m$^7$GpppU, m$^7$GpppG, m$^7$GpppA, m$^7$Gpppm$^6$A, m$^7$GpppCm, m$^7$GpppUm, m$^7$GpppGm, m$^7$GpppAm, m$^7$Gpppm$^6$Am, NAD, FAD, UDP-Glc, UDP-GlcNAc, m$^1$A, m$^6$A, m$^1$Am, m$^6$Am.

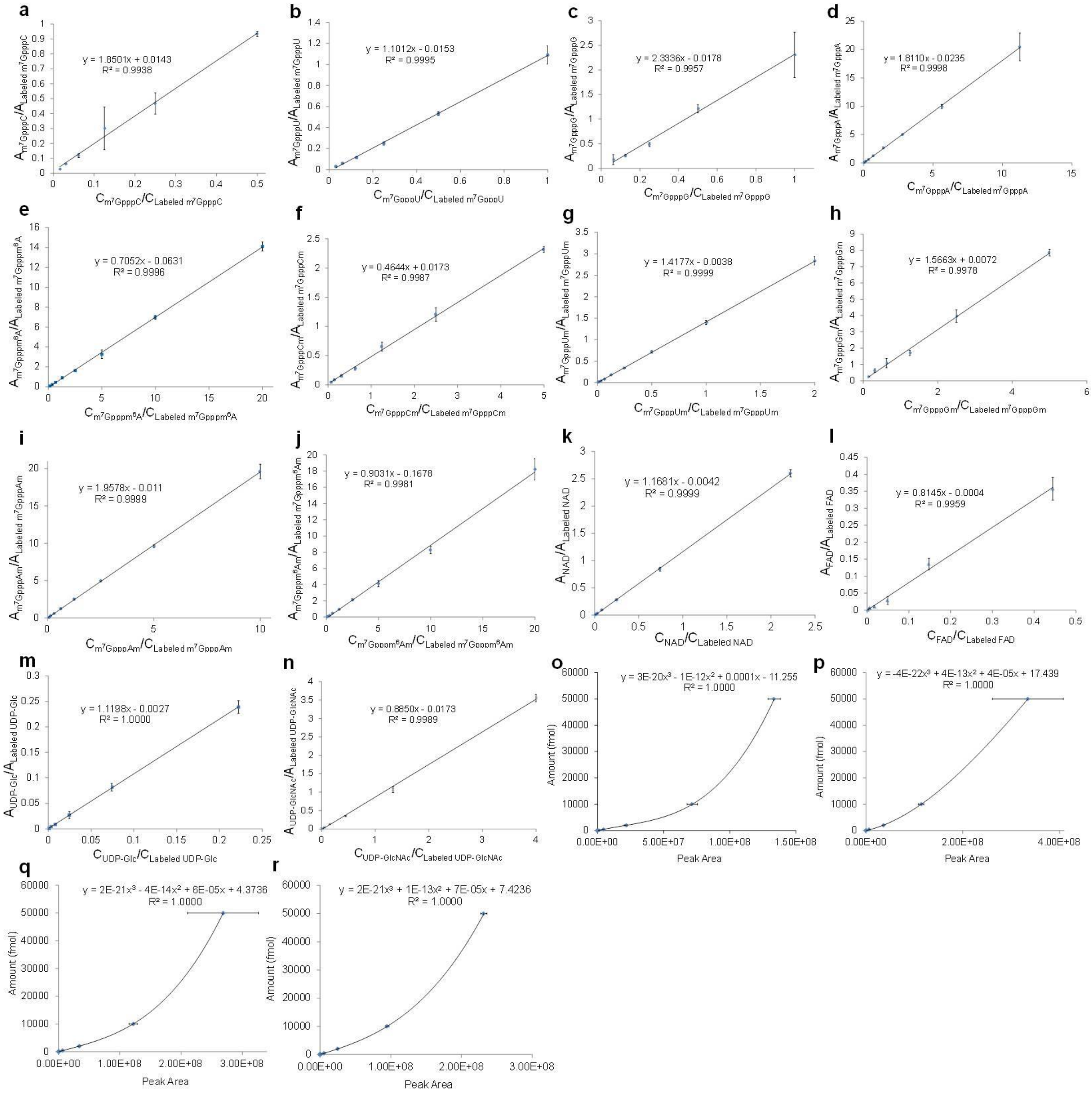

**Figure S10.** Quantification of the release of caps from m$^7$GpppN- and m$^7$GpppNm-capped RNA oligos during NP1 digestion (N = C, U, G, A or m$^6$A). (**a**) m$^7$GpppN-capped RNA oligos; (**b**) m$^7$GpppNm-capped RNA oligos.

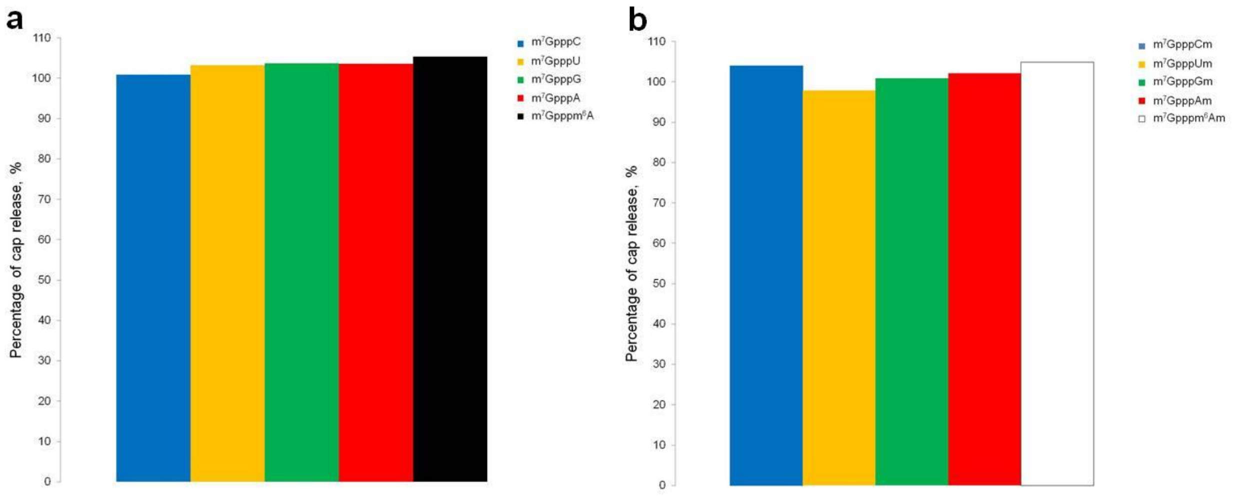

**Figure S11.** Analysis of m$^7$Gpppm$^1$A and m$^7$Gpppm$^1$Am in RNA. (**a**) Off-line HPLC enrichment: a representative trace for the off-line HPLC enrichment of m$^7$Gpppm$^1$A and m$^7$Gpppm$^1$Am from the enzymatic digestion mixture of cellular RNA. (**b**) LC-MS/MS analysis: HPLC elution profiles and MS/MS transitions (X→Y) for m$^7$Gpppm$^1$A and m$^7$Gpppm$^1$Am in human CCRF-SB mRNA.

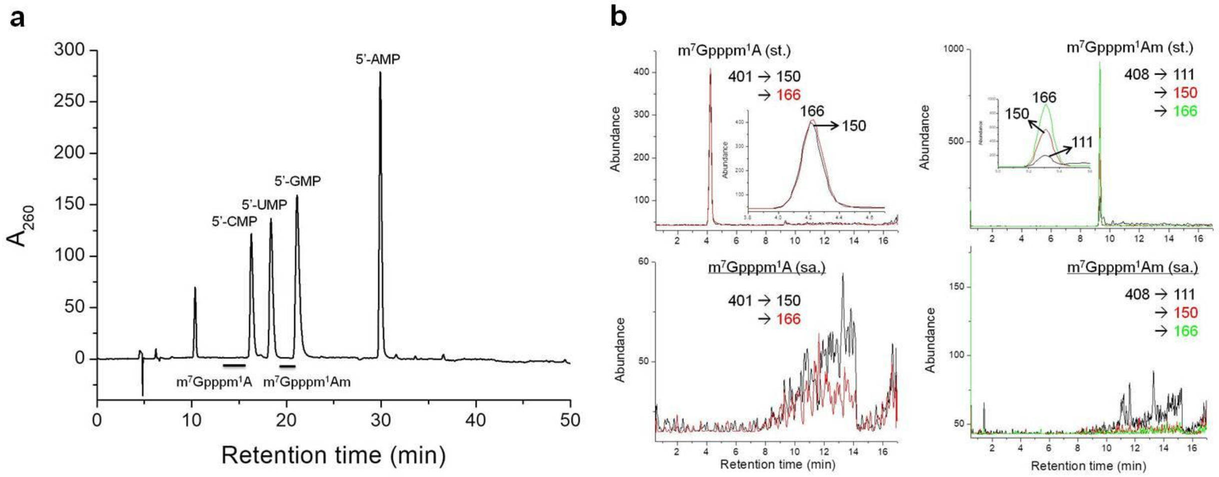

**Figure S12.** Quantitative real-time PCR analysis of the expression of CMTR1, PCIF1 FTO, DCP2 and ALKBH5 genes in CCRF-SB, mouse liver, and mouse kidney total RNA. Error bars represent mean ± s.d. for n = 3 biological replicates.

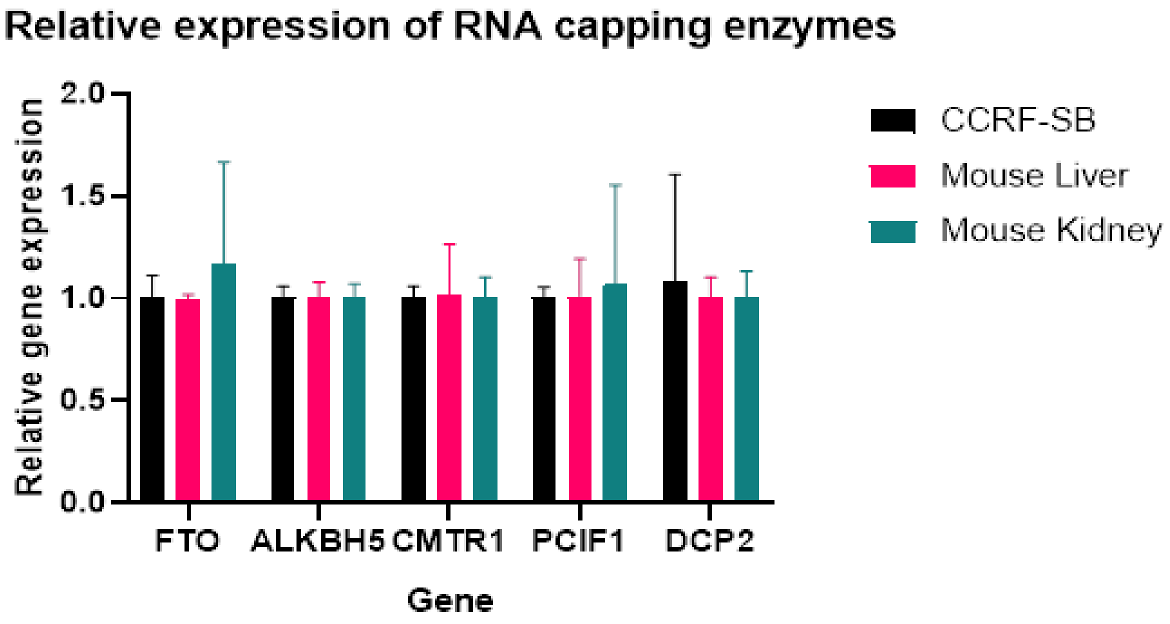

**Table S1.** Detection limits of known and potentially existing cap nucleotides studied.

| Compound | Limit of quantification (LOQ), fmol | Limit of detection (LOD), fmol |
|---|---|---|
| $m^7$GpppC | 6.1 | 1.8 |
| $m^7$GpppU | 0.30 | 0.09 |
| $m^7$GpppG | 25 | 7.5 |
| $m^7$GpppA | 7.3 | 2.2 |
| $m^7$Gpppm$^6$A | 3.1 | 0.93 |
| $m^7$GpppCm | 9.7 | 3.0 |
| $m^7$GpppUm | 0.062 | 0.019 |
| $m^7$GpppGm | 41 | 12 |
| $m^7$GpppAm | 2.0 | 0.60 |
| $m^7$Gpppm$^6$Am | 4.0 | 1.2 |
| GpppC | 240 | 72 |
| GpppU | 4.3 | 1.3 |
| GpppG | 38 | 11 |
| GpppA | 8.7 | 2.6 |
| Gpppm6A | 43 | 13 |
| GpppCm | 540 | 160 |
| GpppUm | 6.9 | 2.1 |
| GpppGm | 380 | 110 |
| GpppAm | 40 | 12 |
| Gpppm$^6$Am | 6.4 | 1.9 |
| NAD | 2.4 | 0.72 |
| FAD | 2.2 | 0.66 |
| UDP-Glc | 0.36 | 0.11 |
| UDP-GlcNAc | 3.1 | 0.93 |
| dpCoA | 7.7 | 2.3 |
| $m^{2,2,7}$GpppG | 6.4 | 1.9 |
| $m^7$Gpppm$^1$A | 2.3 | 0.68 |
| $m^7$Gpppm$^1$Am | 0.38 | 0.11 |

**Table S2.** Exact mass of synthetic unlabeled cap dinucleotides determined by high-resolution mass spectrometry.

| Cap dinucleotide | Molecular formula | $[M+H]^+$ calculated | $[M+H]^+$ measured | $\Delta_m$, ppm |
|---|---|---|---|---|
| GpppC | $C_{19}H_{27}N_8O_{18}P_3$ | 749.0735 | 749.0741 | 0.80 |
| GpppU | $C_{19}H_{26}N_7O_{19}P_3$ | 750.0575 | 750.0539 | 4.8 |
| Gpppm$^6$A | $C_{21}H_{29}N_{10}O_{17}P_3$ | 787.1003 | 787.1021 | 2.3 |
| GpppC$_m$ | $C_{20}H_{29}N_8O_{18}P_3$ | 763.0891 | 763.0862 | 3.8 |
| GpppU$_m$ | $C_{20}H_{28}N_7O_{19}P_3$ | 764.0731 | 764.0754 | 3.0 |
| GpppA$_m$ | $C_{21}H_{29}N_{10}O_{17}P_3$ | 787.1003 | 787.1001 | 0.25 |
| GpppG$_m$ | $C_{21}H_{29}N_{10}O_{18}P_3$ | 803.0952 | 803.0927 | 3.1 |
| Gpppm$^6$A$_m$ | $C_{22}H_{31}N_{10}O_{17}P_3$ | 801.1160 | 801.1145 | 1.9 |
| m$^7$GpppC | $C_{20}H_{29}N_8O_{18}P_3$ | 763.0891 | 763.0924 | 4.3 |
| m$^7$GpppU | $C_{20}H_{28}N_7O_{19}P_3$ | 764.0731 | 764.0750 | 2.5 |
| m$^7$Gpppm$^6$A | $C_{22}H_{31}N_{10}O_{17}P_3$ | 801.1160 | 801.1169 | 1.1 |
| m$^7$GpppC$_m$ | $C_{21}H_{31}N_8O_{18}P_3$ | 777.1048 | 777.1071 | 3.0 |
| m$^7$GpppU$_m$ | $C_{21}H_{30}N_7O_{19}P_3$ | 778.0888 | 778.0899 | 1.4 |
| m$^7$GpppA$_m$ | $C_{22}H_{31}N_{10}O_{17}P_3$ | 801.1160 | 801.1162 | 0.25 |
| m$^7$GpppG$_m$ | $C_{22}H_{31}N_{10}O_{18}P_3$ | 817.1109 | 817.1100 | 1.1 |
| m$^7$Gpppm$^6$A$_m$ | $C_{23}H_{33}N_{10}O_{17}P_3$ | 815.1316 | 815.1334 | 2.2 |

**Table S3.** Primers used for qPCR analysis. Primer pairs span across exon-exon junctions and separated by intron where possible.

| | | |
|---|---|---|
| **Gapdh Human qPCR Primer Pair (NM_002046)** | | |
| F | GTCTCCTCTGACTTCAACAGCG | product length = 131 |
| R | ACCACCCTGTTGCTGTAGCCAA | |
| **POLR2A Human qPCR Primer Pair (NM_000937)** | | |
| F | GAGAGCGTTGAGTTCCAGAACC | product length = 152 |
| R | TGGATGTGTGCGTTGCTCAGCA | |
| **FTO Human qPCR Primer Pair (NM_001080432)** | | |
| F | CCAGAACCTGAGGAGAGAATGG | product length = 143 |
| R | CGATGTCTGTGAGGTCAAACGG | |
| **ALKBH5 Human qPCR Primer Pair (NM_017758)** | | |
| F | CCAGCTATGCTTCAGATCGCCT | product length = 132 |
| R | GGTTCTCTTCCTTGTCCATCTCC | |
| **CMTR1 Human qPCR Primer Pair (NM_015050)** | | |
| F | GCTGCTTCTGTGTCAGTTCCTC | product length = 120 |
| R | CAGTACAGCAGGTAGACAAGCC | |
| **PCIF1 Human qPCR Primer Pair (NM_022104)** | | |
| F | CTCTGCCTTTGAGAGGTTCCTG | product length = 161 |
| R | AGCACTCGAAGCTGACGCCAAA | |
| **DCP2 Human qPCR Primer Pair (NM_152624)** | | |
| F | AGCAGCAGAATTCTTTGATGAAGTG | product length = 108 |
| R | GGTCTTCACTGGAGCTAGGC | |
| **Gapdh Mouse qPCR Primer Pair (NM_008084)** | | |
| F | AGGTCGGTGTGAACGGATTTG | product length = 123 |
| R | TGTAGACCATGTAGTTGAGGTCA | |
| **Polr2a Mouse qPCR Primer Pair (NM_009089)** | | |
| F | CGAGAAGGTCTCATTGACACGG | product length = 131 |
| R | ACCACCTGGTTGATGGAGTTCC | |
| **Fto Mouse qPCR Primer Pair (NM_011936)** | | |
| F | GCCTCGGTTTAGTTCCACTCAC | product length = 118 |
| R | GTCGCCATCGTCTGAGTCATTG | |
| **Alkbh5 Mouse qPCR Primer Pair (NM_172943)** | | |
| F | GTGGGTATGCTGCTGATGAAATC | product length = 108 |
| R | CCAATCGCGGTGCATCTAATC | |
| **Cmtr1 Mouse qPCR Primer Pair (NM_028791)** | | |
| F | CCATCCCCTCAGCTAACAATGAA | product length = 173 |
| R | CCACTGAGGCTTGTAGCAGAT | |
| **Pcif1 Mouse qPCR Primer Pair (NM_146129)** | | |
| F | CTCTGCCTTTGAGAGGTTCCTG | product length = 155 |
| R | CGAAACTGACACCAAAGAGTCGG | |
| **Dcp2 Mouse qPCR Primer Pair (NM_027490)** | | |

| | | |
|---|---|---|
| F | GCAGAATCCCTTGGTGAAATGTG | product length = 109 |
| R | CGACGGCTCTTCACCAGAAC | |
